# Evidence for a Compensatory Relationship between Left- and Right-Lateralized Brain Networks

**DOI:** 10.1101/2023.12.08.570817

**Authors:** Madeline Peterson, Rodrigo M. Braga, Dorothea L. Floris, Jared A. Nielsen

## Abstract

The two hemispheres of the human brain are functionally asymmetric. At the network level, the language network exhibits left-hemisphere lateralization. While this asymmetry is widely replicated, the extent to which other functional networks demonstrate lateralization remains a subject of investigation. Additionally, it is unknown how the lateralization of one functional network may affect the lateralization of other networks within individuals. We quantified lateralization for each of 17 networks by computing the relative surface area on the left and right cerebral hemispheres. After examining the ecological, convergent, and external validity and test-retest reliability of this surface area-based measure of lateralization, we addressed two hypotheses across multiple datasets (Human Connectome Project = 553, Human Connectome Project-Development = 343, Natural Scenes Dataset = 8). First, we hypothesized that networks associated with language, visuospatial attention, and executive control would show the greatest lateralization. Second, we hypothesized that relationships between lateralized networks would follow a dependent relationship such that greater left-lateralization of a network would be associated with greater right-lateralization of a different network within individuals, and that this pattern would be systematic across individuals. A language network was among the three networks identified as being significantly left-lateralized, and attention and executive control networks were among the five networks identified as being significantly right-lateralized. Next, correlation matrices, an exploratory factor analysis, and confirmatory factor analyses were used to test the second hypothesis and examine the organization of lateralized networks. We found general support for a dependent relationship between highly left-and right-lateralized networks, meaning that across subjects, greater left lateralization of a given network (such as a language network) was linked to greater right lateralization of another network (such as a ventral attention/salience network) and vice versa. These results further our understanding of brain organization at the macro-scale network level in individuals, carrying specific relevance for neurodevelopmental conditions characterized by disruptions in lateralization such as autism and schizophrenia.

## 1 Introduction

Observations of the human brain have revealed significant differences in the gross anatomical morphometry between the two hemispheres (for review, see (Toga & Thompson, 2003). These structural asymmetries are accompanied by functional asymmetries, most notably for language specialization. Famously, Paul Broca localized language specialization to the left hemisphere subsequent to identifying a lesion in the left inferior frontal gyrus of his patient as being responsible for his eponymous aphasia (Broca, 1861). This contribution launched an emphasis on regions specialized for language, which were later conceptualized as a network consisting of Broca’s and Wernicke’s areas connected via the arcuate fasciculus (Geschwind, 1972).

Contemporarily, the language network is regarded as a prototypical example of a lateralized network, with left-hemisphere language lateralization estimated to occur in most (Breier et al., 2000; Stippich et al., 2003) to more than 90% of the general population (Corballis, 2003). The canonical language network is a distributed network comprising regions across the frontal, temporal, and parietal lobes, with lines of evidence stemming from a variety of sources including lesion cases (Broca, 1861; Geschwind, 1970; Wernicke, 1874), intraoperative brain stimulation (Penfield & Jasper, 1954), neurodegeneration (e.g., primary progressive aphasia; Mesulam, 2001, 2003; Mesulam et al., 2014, 2015), task-based fMRI (Fedorenko et al., 2010, 2011; Fedorenko, McDermott, et al., 2012, 2012; Lipkin et al., 2022; Scott et al., 2017), and functional connectivity (Braga et al., 2020; Hacker et al., 2013; Lee et al., 2012). The typically asymmetric organization of this network in neurotypical individuals continues to be replicated (Elin et al., 2022; Malik-Moraleda et al., 2022; Olulade et al., 2020; Reynolds et al., 2019).

While the lateralization of language provides a compelling example, it also prompts broader questions about the origins and implications of cerebral lateralization across other cognitive domains. In attempting to unravel the origins of cerebral lateralization, researchers have explored theoretical perspectives ranging from the genetic and epigenetic (Geschwind & Miller, 2001; McManus, 1985) to interhemispheric conflict (Andrew et al., 1982; Corballis, 1991). Yet, these paradigms fall short in explaining the dynamic interactions of interdigitated lateralized and non-lateralized networks. For example, it is unclear how having a highly lateralized network, such as the language network, may influence the lateralization of other networks within individuals. Along the lines of the interhemispheric conflict explanation of lateralization, competition for limited cortical resources during brain maturation may drive lateralization. According to this hypothesis, as different functional networks vie for cortical real estate and resources, they become lateralized. Alternatively, or in tandem with this mechanism, networks may become lateralized in order to optimize their efficiency, preventing interference from competing networks. Under this framework, as one network increases in lateralization to one hemisphere, that network occupies more space within that hemisphere while freeing up cortical territory in the contralateral hemisphere. Presumably, this would allow for a complimentary network to become more lateralized in the contralateral hemisphere. One example of such a scenario may be found in a right-lateralized attention network composed of the temporoparietal junction and ventral frontal areas and which is hypothesized to process visuospatial information, particularly unexpected stimuli (Corbetta & Shulman, 2002). The ventral attention network in particular has been identified as a potential right-lateralized compliment to the left-lateralized language network (Bernard et al., 2020).

Other functional networks in both the right and left hemispheres have been examined for evidence of lateralization. Of note, lateralization can be an indicator for specialization, or the dominant hosting of a macroscale functional network and its associated functional properties by one hemisphere over the other (Hervé et al., 2013). One study quantified specialization across seven functional networks and found that specialization was not restricted to a single left-or right-specialized network (Wang et al., 2014). Rather, the right frontoparietal network and right ventral and dorsal attention networks, as well as the left default and frontoparietal networks exhibited specialization as assessed via a functional connectivity-based metric (see Fig. 5; Wang et al., 2014). This pattern was generally replicated in highly sampled individuals, revealing left-lateralized language, default, and frontoparietal networks, as well as right-lateralized salience and frontoparietal networks (Braga et al., 2020). Interestingly, the finding of both left-and right-lateralized frontoparietal networks across both Braga et al. (2020) and Wang et al. (2014) evidences a joint control system in which a subdivision of the frontoparietal control network is coupled with other lateralized networks in either the left or right hemisphere. Beyond this result, research on network lateralization has untapped potential when it comes to understanding the relationships between lateralized networks. This includes associations in laterality between ipsilateral and contralateral lateralized networks and extends to patterns within and across individuals.

### 1.1 Methods for Examining Hemispheric Asymmetries

In humans, hemispheric specialization has historically been identified using a variety of methods including callosotomy (i.e., split-brain patients; for review, see Gazzaniga, 2000), lateralized brain lesions (Milner, 1971; Rasmussen & Milner, 1977), the unilateral carotid administration of anesthetic (i.e., the Wada test Wada & Rasmussen, 1960), and intraoperative brain stimulation mapping (Penfield & Jasper, 1954). Callosotomy studies have revealed the importance of interhemispheric communication for certain cognitive processes, demonstrating that the left and right hemispheres can operate relatively independently for some functions but require communication for others (Gazzaniga, 2000). Lateralized brain lesion studies, particularly the work of Milner and Rasmussen, have identified specific functions associated with each hemisphere, such as language processing predominantly in the left hemisphere (Milner, 1971; Rasmussen & Milner, 1977). Similarly, the Wada test has shed light on hemispheric dominance for language and memory (Wada & Rasmussen, 1960). Finally, leaning into the localization of specific functions to certain regions within each hemisphere, intraoperative brain stimulation mapping has provided detailed maps of functional areas in the brain (Penfield & Jasper, 1954). Collectively, these classic methods reveal patterns of human brain organization governed by interactions between lateralization and localization.

These historical methods are complimented by neuroimaging metrics, many of which are functional connectivity-based. Of particular interest are the intrinsic laterality index (Liu et al., 2009), autonomy index (Wang et al., 2014), hemispheric contrast (Gotts et al., 2013), functional lateralization metric (Nielsen et al., 2013), classification metric (Friedrich et al., 2022), and network variants approach (Perez et al., 2023). Despite the unifying aim of estimating hemispheric specialization or lateralization, each of the listed methods varies in terms of how it approaches structural asymmetries, the addition of covariates such as handedness and gender, and short-and long-range connectivity. However, with the exception of the network variants approach (Perez et al., 2023), each method has been implemented on less than 12 minutes of resting-state fMRI data per participant, a tactic which is increasingly being exchanged for a within-individual “precision” approach.

### 1.2 A Precision Approach to Lateralization

The precision approach, which emphasizes extensive individual sampling, is being heralded as a well-powered alternative to the large and costly sample sizes required for cross-sectional group and brain-wide association studies (Gratton et al., 2022; Marek et al., 2022). This method of densely sampling individuals can generate precise brain maps (Gordon et al., 2017) as well as the development of optimal workflows for reducing MRI artifacts (Ciric et al., 2017). Moreover, when combined with functional localizers, the precision approach offers superior sensitivity, functional resolution, and interpretability (Fedorenko, 2021). As applied to estimating lateralization, repeated sampling can improve measures of individual network parcellations (Braga et al., 2020; Gordon et al., 2017, 2020) including network topology and topography, and functional connectivity (Gordon et al., 2017; Laumann et al., 2015), resulting in more precise lateralization measures.

In line with the precision neuroimaging approach and previous efforts to understand brain network organization and lateralization, the present study examines two questions. First, we explore which networks exhibit the greatest hemispheric asymmetries. A recent study involving 18 densely-sampled individuals demonstrated that among six networks, the language network displayed the greatest left hemisphere lateralization, while a frontoparietal control network exhibited the greatest right hemisphere lateralization (Braga et al., 2020). However, it remains unclear how these estimates might change in a larger sample with a greater number of examined networks. Building upon the work of Braga et al. (2020), we hypothesized that networks associated with language, visuospatial attention, and executive control would show the greatest hemispheric asymmetries.

Second, we investigate how lateralization in one network may influence the lateralization of other networks. We propose the following hypotheses to guide our investigation. The first hypothesis suggests that if an individual possesses a highly lateralized network, other networks for that individual will exhibit increased lateralization in the opposite direction, and that this dependent relationship will be systematic across individuals (the dependent hypothesis). The alternative hypothesis proposes that lateralization will be unrelated between networks across individuals (the independent hypothesis).

## 2 Methods

### 2.1 Datasets and Overview

Three independent datasets were used for these analyses: The Human Connectome Project (HCP; split into discovery and replication datasets), the Human Connectome Project-Development (HCPD; Somerville et al., 2018), and the Natural Scenes Dataset (NSD; Allen et al., 2022). Each dataset was selected for its relatively high quantity of low-motion data per participant (see Figure 1).

**Figure 1.**
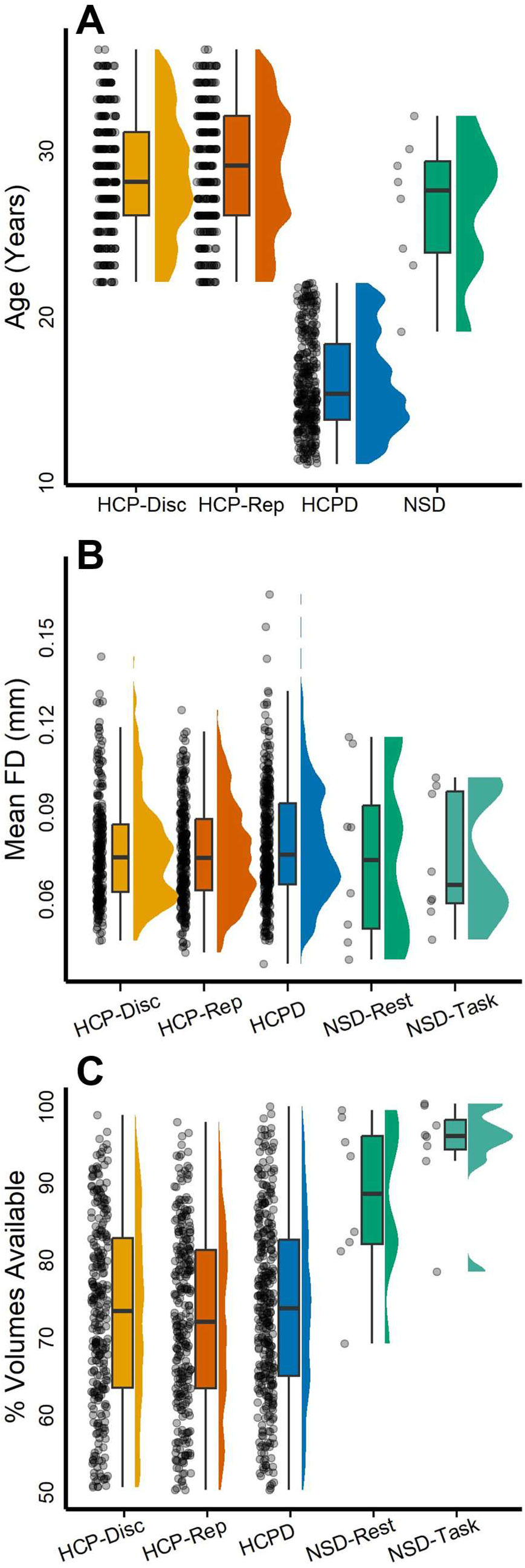
Participant age, data quality, and data availability. Panel A depicts participant age across each dataset following the implementation of exclusion criteria. HCP-Discovery participants included 276 individuals 22-36 years of age, HCP-Replication participants included 277 individuals 22-36 years of age, HCPD participants included 343 individuals 11-22 years of age, and NSD participants included eight individuals 19-32 years of age. Panel B depicts the mean framewise displacement (FD) across each dataset following the implementation of exclusion criteria. HCP-Discovery mean FD was 0.08 mm (*SD* = 0.02 mm), range 0.04-0.14 mm; HCP-Replication mean FD was 0.07 mm (*SD* = 0.01 mm), range 0.04-0.12 mm; HCPD mean FD was 0.08 mm (*SD* = 0.02 mm), range 0.04-0.16 mm; NSD-Rest mean FD was 0.07 mm (*SD* = 0.03), range 0.04-0.11 mm; and NSD-Task mean FD was 0.07 mm (*SD* = 0.02 mm), range 0.04-0.1 mm. Panel C depicts the percentage of volumes remaining following motion-correction procedures for each dataset. HCP-Discovery mean percentage of volumes was 72.81% (*SD* = 12.12%), range 50.38-98.54%; HCP-Replication mean percentage of volumes was 72.09% (*SD* = 11.59%), range 50.04-97.62%; HCPD mean percentage of volumes was 73.6% (*SD* = 11.73%), range 50.05-99.63%; NSD-Rest mean percentage of volumes was 87.64% (*SD* = 10.51%), range 68.98-99.14%; and NSD-Task mean percentage of volumes was 94.27% (*SD* = 6.92), range 78.31-100%. Across each panel, a circle represents a single participant.

#### 2.1.1 HCP Discovery and Replication

The HCP S1200 release consists of 1206 subjects (1113 with structural MRI scans) collected at 13 different data acquisition sites with informed consent (Van Essen et al., 2013). Additional details regarding HCP scanning protocols are available online (https://humanconnectome.org/storage/app/media/documentation/s1200/HCP_S1200_Release_A ppendix_I.pdf; Uğurbil et al., 2013; Van Essen et al., 2012). With a relatively large quantity of data available per individual, this data is ideally suited for taking a within-individual approach to specialization. Participants underwent four 15-minutes runs of a passive fixation task (resting-state fMRI) during which they were asked to keep their eyes open while viewing a white cross on a dark background and think of nothing in particular while remaining awake (Smith et al., 2013). Exclusion criteria for the HCP S1200 release included removing participants with a mean framewise displacement greater than 0.2 mm and mean DVARS greater than 50, participants missing handedness data, and participants with less than 50% of volumes remaining after motion censoring. This resulted in a subsample of 553 participants, which was split into a discovery and replication dataset using random sampling without replacement. The two datasets were then compared using the R package MatchIt (Ho et al., 2023) on age, mean framewise displacement, sex, handedness, and percentage of volumes remaining following motion censoring. The HCP-Discovery dataset consisted of 276 participants 22-36 years old (*M* = 28.48, *SD* = 3.58) with 167 females, while the HCP-Replication dataset consisted of 277 participants 22-36 years old (*M* = 28.7, *SD* = 3.77) with 173 females.

#### 2.1.2 HCPD

With a younger sample and smaller quantity of data per individual, the HCPD dataset was used as an additional replication dataset for primary analyses. Since data collection for the HCPD project is ongoing, cross-sectional data from the latest release were included, and these were composed of 652 healthy participants. All data were obtained with informed assent or consent. As a part of the HCPD protocol, participants underwent four 6.5-minute runs of resting-state fMRI, with an exception for participants 5-7 years old, which had six 3.5-minute runs each (Harms et al., 2018). Participants were instructed to view a small white fixation crosshair on a black background and blink normally. Exclusion criteria for HCPD included removing participants with less than 50% of volumes remaining after motion censoring, participants missing handedness data, and participants with a mean framewise displacement greater than 0.2 mm and mean DVARS greater than 50 (see Figure 1). Following the exclusion criteria, the dataset consisted of 343 individuals ages 11-21.92 (*M* = 15.93, *SD* = 2.97) of which 189 were female.

#### 2.1.3 NSD

With a large quantity of resting-state and task fMRI data available per individual, the NSD was included to examine potential task effects on estimating individual network parcellations and specialization. The NSD is composed of eight individuals (two males and six females; age range 19–32 years). All data were obtained with informed written consent according to the University of Minnesota institutional review board. As detailed in Allen et al. (2022), participants averaged two hours of resting state fMRI and 39.5 hours of task-based fMRI. For the resting-state runs, participants were instructed to stay awake and fixate on a white cross placed on a gray background but otherwise rest. During the task-based runs, participants were shown distinct natural scenes taken from the Microsoft Common Objects in Context database (T.-Y. Lin et al., 2014). Images were presented for 3 s with 1-s gaps in between images. Subjects fixated centrally and performed a long-term continuous recognition task on the images. Exclusion criteria for NSD included removing participants with less than 50% of volumes remaining after motion censoring, and participants with a mean framewise displacement greater than 0.2 mm and mean DVARS greater than 50. No subjects were excluded from the analysis; however, following motion correction, a minimum of 12 resting-state fMRI runs (approximately 60 minutes) remained. In order to compare resting-state and task data on equal grounds, only the first 12 available resting-state runs and the first 12 available task fMRI runs from each participant were utilized.

### 2.2 MRI Acquisition Parameters

#### 2.2.1 HCP Discovery and Replication

The HCP dataset was acquired on a custom Siemens 3T Skyra with a 32-channel head coil. T1-weighted images were collected with a 3D MPRAGE sequence with isotropic 0.7 mm voxels (256 sagittal slices, repetition time [TR] = 2400 milliseconds, echo time [TE] = 2.14 milliseconds) as detailed in Glasser et al. (2013). Resting-state functional images were collected using 2 mm isotropic voxels (72 sagittal slices, TR = 720 milliseconds, TE = 33 milliseconds, multiband accelerated pulse sequence with multiband factor = 8) as detailed in Glasser et al. (2013, 2016).

#### 2.2.2 HCPD

The HCPD MRI data were acquired on Siemens 3T Prisma scanners with vendor 32-channel headcoils at four sites: Harvard University, University of California-Los Angeles, University of Minnesota, and Washington University in St. Louis (Harms et al., 2018). Structural T1-weighted scans were acquired with a multi-echo MPRAGE sequence (van der Kouwe et al., 2008) with 0.8 mm isotropic voxels (sagittal FOV = 256 × 240 × 166; matrix size = 320 × 300 × 208 slices; slice oversampling = 7.7%; 2-fold in-plane acceleration (GRAPPA); pixel bandwidth = 744 Hz/Px; Tr/TI = 2500/1000, TE = 1.9/3.6/5.4/7.2 ms, flip angle = 8°; water excitation employed for fat suppression; up to 30 TRs allowed for motion-induced reacquisition). T2*-weighted scans were used for resting-state fMRI with 2D multiband gradient-recalled echo echo-planar imaging sequence (MB8, TR/TE = 800/37 ms, flip angle = 52°) and 2.0 mm isotropic voxels covering the whole brain (72 oblique-axial slices). Functional scans were acquired in pairs of two runs with opposite phase encoding polarity (anterior-to-posterior and posterior-to-anterior) so that fMRI data were not biased towards either phase encoding polarity. For all scans, Framewise Integrated Real-time MRI Monitoring (Dosenbach et al., 2017) was implemented to provide motion feedback to participants between fMRI runs.

#### 2.2.1 NSD

The NSD dataset was acquired at the Center for Magnetic Resonance Research at the University of Minnesota (Allen et al., 2022). Anatomical data (such as T1-weighted volumes) were collected using a 3T Siemens Prisma scanner with a standard Siemens 32-channel RF head coil while functional data were collected using a 7T Siemens Magnetom passively shielded scanner and a single-channel-transmit, 32-channel-receive RF head coil. T1-weighted images were acquired with a MPRAGE sequence (0.8-mm bandwidth 220 Hz per pixel, no partial Fourier, in-plane acceleration factor (iPAT) 2, TA = 6.6 min per scan). Functional data were collected using gradient-echo EPI at 1.8-mm isotropic resolution with whole-brain coverage (84 axial slices, slice thickness 1.8 mm, slice gap 0 mm, field-of-view 216 mm (FE) *×* 216 mm (PE), phase encode direction anterior-to-posterior, matrix size 120 *×* 120, TR = 1,600 milliseconds, TE = 22.0 milliseconds, flip angle 62°, echo spacing 0.66 milliseconds, bandwidth 1,736 Hz per pixel, partial Fourier 7/8, iPAT 2, multi-band slice acceleration factor 3). Full protocol printouts for the NSD dataset are available online (https://cvnlab.slite.page/p/NKalgWd F/Experiments).

### 2.3 fMRI Preprocessing

Preprocessing took place on raw NIFTI files for the resting-state fMRI and task fMRI runs using a pipeline developed by the Computational Brain Imaging Group (CBIG; Kong et al., 2019; Li et al., 2019; code is available online at https://github.com/ThomasYeoLab/CBIG/tree/c773720ad340dcb1d566b0b8de68b6acdf2ca505/s table_projects/preprocessing/CBIG_fMRI_Preproc2016). This CBIG2016 preprocessing pipeline was selected to process the fMRI data in order to more closely follow the processing steps used to implement the multi-session hierarchical Bayesian modeling parcellation method (Kong et al., 2019). As a prerequisite, this pipeline requires FreeSurfer recon-all output from the structural data (FreeSurfer 6.0.1; Dale et al., 1999). The fMRI data are then processed with the following steps: 1) removal of the first four frames and 2) motion correction using rigid body translation and rotation with the FSL package (Jenkinson et al., 2002; Smith et al., 2004). The structural and functional images are then aligned using boundary-based registration (Greve & Fischl, 2009) using the FsFast software package (http://surfer.nmr.mgh.harvard.edu/fswiki/FsFast). FD and DVARS were computed using *fsl_motion_outliers* (Smith et al., 2004). Volumes with FD > 0.2 mm or DVARS > 50 were tagged as outliers. Uncensored segments of data lasting fewer than 5 contiguous volumes were also flagged as outliers (Gordon et al., 2016). BOLD runs with more than half of the volumes flagged as outliers were removed completely. Next, linear regression using multiple nuisance regressors was applied through a combination of CBIG in-house scripts and the FSL MCFLIRT tool (Jenkinson et al., 2002). Nuisance regressors consisted of global signal, six motion correction parameters, averaged ventricular signal, averaged white matter signal, and their temporal derivatives (totaling 18 regressors). The flagged outlier volumes were ignored during the regression procedure. Following the regression, a bandpass filter (0.009 Hz ≤ f ≤ 0.08 Hz) was applied using CBIG in-house scripts. At this point, the preprocessed fMRI data were projected onto the FreeSurfer fsaverage6 surface space (2 mm vertex spacing) with FreeSurfer’s *mri_vol2surf* function. The projected fMRI data were then smoothed using a 6 mm full-width half-maximum kernel through FreeSurfer’s *mri_surf2surf* function (Fischl et al., 1999). Surface space was selected for the following analyses in order to best follow the individual parcellation pipeline outlined in Kong et al. (2019), and following evidence that landmark surface-based registration outperforms volume-based registration (Anticevic et al., 2008; Argall et al., 2006; Desai et al., 2005; Van Essen, 2005).

### 2.4. Individual Network Parcellation

Following preprocessing, network parcellations were computed using a multi-session hierarchical Bayesian modeling (MS-HBM) pipeline. The MS-HBM pipeline is designed to generate parcellations for individuals with multiple sessions of fMRI data (Kong et al., 2019; Li et al., 2019) and is implemented in MATLAB R2018b (MATLAB, 2018). This particular model has been selected because it accounts for intra-individual variation, allowing the model to better generalize to new fMRI data from the same participant. As an overview, this model uses a variational Bayes expectation-maximization algorithm to learn group-level priors from a training dataset and then apply those to estimate individual-specific parcellations (see Figure 2). This model estimates the following parameters: group-level network connectivity profiles, inter-subject functional connectivity variability, intra-subject functional connectivity variability, a spatial smoothness prior, and an inter-subject spatial variability prior. As recommended in the pipeline’s GitHub documentation, subjects with a single available run post-preprocessing had that single run split in two and a connectivity profile was generated for each split. A *k* of 17 was selected for all participants (Yeo et al., 2011). Additionally, it has previously been demonstrated that MS-HBM parameters estimated from one dataset can be effectively applied to another dataset with significant differences in acquisition and preprocessing (Kong et al., 2021). Thus, to generate our model, priors trained on 37 Genomic Superstruct Project (GSP) subjects were utilized (Holmes et al., 2015; Kong et al., 2019). Following the generation of individual parcellations, a Hungarian matching algorithm was used to match the clusters with the Yeo et al. (2011) 17-network group parcellation.

**Figure 2.**
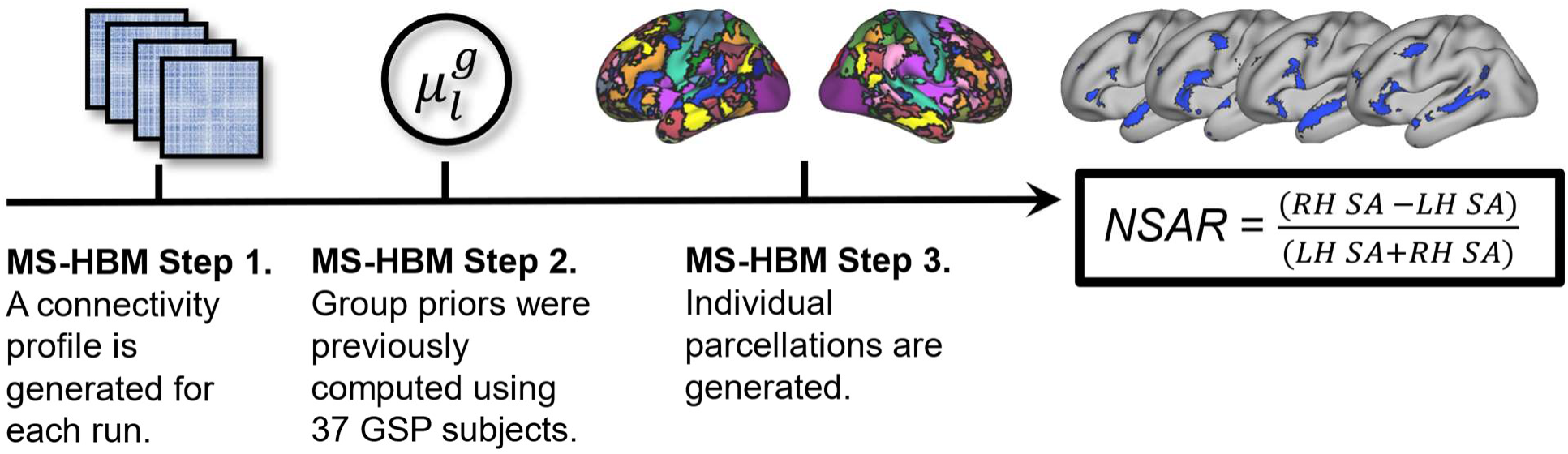
Illustration of the Multi-Session Hierarchical Bayesian Modeling (MS-HBM) individual parcellation pipeline. First, a connectivity profile is generated for each available fMRI run on an individual basis (illustrated here as a functional connectivity matrix). Next, group priors previously estimated (Kong et al., 2019) from 37 Genomic Superstruct Project (GSP) subjects were used. Third, the connectivity profiles from each available run and the group priors (more specifically, the inter-subject functional connectivity variability, intra-subject functional connectivity variability, spatial smoothness, and inter-subject spatial variability) are used to generate network parcellations for each participant. Finally, the network surface area ratio (NSAR) is calculated using the formula shown, where LH SA is the left hemisphere surface area for a given network and RH SA is the right hemisphere surface area for a given network. A negative NSAR value indicates left hemisphere lateralization for a given network while a positive value indicates right hemisphere lateralization.

### 2.5 Network Surface Area Ratio

Following the generation of individual network parcellations, lateralization was estimated using a novel measure: the network surface area ratio (NSAR). In discussing this measure, we opted to use this terminology (lateralization) because it accurately encapsulates the concept of an asymmetrical distribution of functional networks across the cerebral hemispheres, which is central to the following analyses. This measure was calculated within each individual for each of 17 networks by first extracting each network label as a region of interest using the Connectome Workbench wb_command functions *metric-label-import* and *gifti-label-to-roi* (Marcus et al., 2013). Next, the left and right hemisphere surface areas for a given network were calculated on a midthickness Conte69 surface in fsaverage6 resolution (Glasser & Essen, 2011) using the wb_command function *metric-stats*. Finally, NSAR was calculated as the difference between normalized left and right hemisphere surface areas for a given network (see Figure 2):

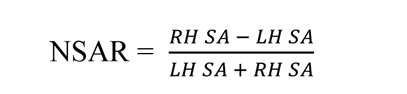

where RH SA represents the right hemisphere surface area of a given network and LH SA represents the left hemisphere surface area of a given network. A scaling factor was not included in the denominator since asymmetry indices including a scaling factor deliver essentially the same findings as those without (Kong et al., 2022).

NSAR values range from -1.0 to +1.0, with negative values indicating left hemisphere lateralization for a given network and positive values indicating right hemisphere lateralization.

NSAR values closer to zero indicate less lateralization (i.e., hemispheric symmetry). Although this measure of lateralization shares similarities with several previously used asymmetry indices (Binder et al., 1997; Braga et al., 2020; Mahowald & Fedorenko, 2016), its distinct methodology prompts us to specifically test its validity and reliability.

### 2.6 Establishing the Validity of NSAR

Ecological validity for Language network laterality was first examined since the Language network has previously been established as a highly lateralized network. HCP subjects with all four runs of resting-state data and the minimally preprocessed Story-Math task contrast were selected (*N* = 221). This Story-Math task was used as a proxy for a language task, as has been done previously (Labache et al., 2023; L. Lin et al., 2022; Wang et al., 2023). Participant *t*-statistic contrast maps were converted to fsaverage6 resolution using wb_command functions *cifti-separate* and *metric-resample*, masked using a language task fMRI atlas (LanA atlas) derived from a large sample (*N* = 804; Lipkin et al., 2022), and then thresholded to the top 10% of vertices. We chose this threshold rather than a fixed *t-*value in order to account for individual differences in the strength of BOLD signal responses attributable to individual differences arising from trait or state factors (Lipkin et al., 2022). A simple laterality metric was then calculated for each contrast map: the number of right hemisphere vertices minus the number of left hemisphere vertices divided by the sum of the left and right hemisphere vertices. A Spearman rank correlation was then used to compare language task laterality against the NSAR value for the Language network. This and all other statistical analyses took place in R 4.2.0 (R Core Team, 2022).

Convergent validity was also examined through a comparison of the NSAR against a measure of specialization: the autonomy index (Wang et al., 2014). The autonomy index approaches specialization from a functional connectivity perspective and is known to reliability estimate specialization across neurotypical and clinical samples (Mueller et al., 2015; Sun et al., 2022; Wang et al., 2014). First, individual functional connectivity matrices were calculated for each resting-state run and then averaged across runs within an individual at the fsaverage6 resolution in MATLAB R2018b (MATLAB, 2018). From here the autonomy index was computed as follows: for each seed ROI obtained from a functional connectivity matrix, the degree of within-hemisphere connectivity and cross-hemisphere connectivity were computed by summing the number of vertices correlated to the seed in the ipsilateral hemisphere and in the contralateral hemisphere. These vertex counts are then normalized by the total number of vertices in the corresponding hemisphere, thus accounting for a potential brain size asymmetry between the two hemispheres. Finally, AI is calculated as the difference between normalized within-and cross-hemisphere connectivity as follows:

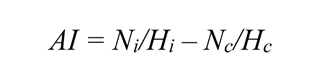

where *N_i_* and *N_c_* are the number of vertices correlated to the seed ROI (using a threshold of |0.25|) in the ipsilateral hemisphere and contralateral hemisphere, respectively. *H_i_* and *H_c_* are the total number of vertices in the ipsilateral and contralateral hemisphere, respectively. To compute the specialization of each functional network, the AI was averaged within the boundary of each network on an individual basis. Subjects from the HCP dataset with all four runs available (*N* = 232) were selected for this analysis of validity and all four runs from each individual were used to compute the autonomy index. A Spearman’s rank correlation coefficient was then used to compare the autonomy index and NSAR on three right-lateralized networks (Limbic-B, Visual-B, and Ventral Attention-A) and three left-lateralized networks (Language, Dorsal Attention-A, and Control-B) determined *a priori*. In order to correct for multiple comparisons, a Bonferroni-corrected alpha level of 0.008 was used.

External validity was next examined through a comparison of NSAR values from significantly lateralized networks against two measures from the Cognition Battery of the National Institutes of Health Toolbox (Gershon et al., 2013): the Oral Reading Recognition Test (ORRT; Gershon et al., 2014) and the Flanker Inhibitory Control and Attention Test (adapted from Rueda et al., 2004). The ORRT was selected as a measure of language and the Flanker as a measure of executive control (inhibitory control, specifically) and visuospatial attention. Among all available cognitive assessments, we selected those that have be shown to engage cognitive domains lateralized to both the right (assessing attention via the Flanker test) and left (evaluating language through the ORRT) hemispheres. Each cognitive measure has been highly validated (Heaton et al., 2014; Ott et al., 2022; Zelazo et al., 2014). To facilitate the comparison of NSAR against these cognitive measures, a Canonical Correlation Analysis (CCA) was implemented using HCP subjects with all four resting-state runs available (*N* = 232). The CCA was chosen for its ability to robustly estimate relationships between sets of variables (Marek et al., 2022), and was conducted using the *cc* function from the CCA package in R (González & Déjean, 2023). CCA feature weights were Haufe-transformed (Haufe et al., 2014) in order to provide a more realistic perspective of feature contributions considering the covariance structure of the data. Haufe-transformations are also known to increase the interpretability and reliability of feature weights (Chen, Ooi, et al., 2022; Chen, Tam, et al., 2022; Tian & Zalesky, 2021)

### 2.7 Establishing the Reliability of NSAR

Reliability analyses sought to address three questions: 1) How much data is needed to obtain a stable estimate of NSAR, 2) What is the test-retest reliability of NSAR, and 3) Is there a task effect on NSAR estimation?

#### 2.7.1 Stable Estimate Analysis

Given that MRI scanning is costly, rendering it comparatively rare to have highly sampled individuals, it is important to understand how much data is needed to reliably estimate lateralization and assess the credibility of our results. To address this concern, we analyzed HCP participants with all four runs of resting-state data available (*N* = 232). Following preprocessing, the first and third scans from each participant were set aside to compose 30 minutes of independent data. Next, the second and fourth scans were each split into three five-minute segments. Runs were split in MATLAB R2018b (MATLAB, 2018) using native MATLAB functions as well as the FreeSurfer functions *MRIread* and *MRIwrite*. The MS-HBM pipeline was then used to generate individual parcellations from 5, 10, 15, 20, 25, and 30 minutes of data from the segmented scans. The MS-HBM pipeline was also used to generate separate individual parcellations from 30 minutes of independent data. Of note, the reliability of the MS-HBM pipeline has been examined previously (see Kong et al., 2019 Figure 3B and Supplementary Figure S10C). The NSAR was then calculated for each iteration (5, 10, 15, etc. minutes) and the independent 30 minutes of data. An intraclass correlation between the NSAR from each iteration parcellation and the independent 30 minutes parcellation was assessed within each subject.

**Figure 3.**
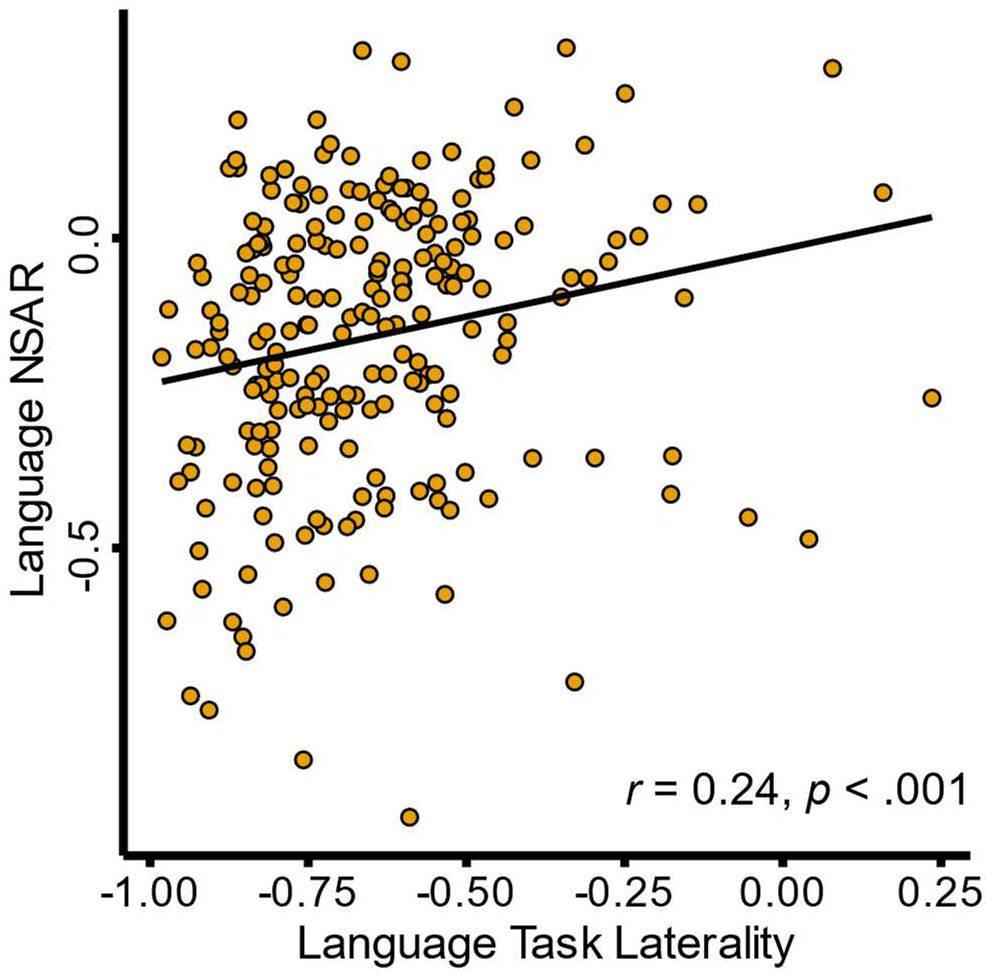
Language network NSAR and language task laterality. Depicted is a positive relationship between NSAR for the Language network and language task laterality in a subset of the HCP dataset (*N* = 221). Across each measure of laterality, a negative value denotes left-hemisphere lateralization while a positive value indicates right-hemisphere lateralization.

Similarly, an intraclass correlation between the NSAR from each iteration parcellation and the independent 30 minutes parcellation was assessed for each network. For the NSAR and parcellation stable estimate analyses, the standard guidelines from Koo & Li (2016) regarding intraclass correlation values were implemented, with values less than 0.5 indicating poor reliability, values between 0.5 and 0.75 indicating moderate reliability, values between 0.75 and 0.9 indicating good reliability, and values greater than 0.9 indicating excellent reliability (based on a 95% confidence interval).

#### 2.7.2 Test-Retest Reliability Analysis

The purpose of the test-retest reliability analysis is to measure the reliability of NSAR in a simpler fashion than the stable estimate analysis. For this analysis, the first two and second two runs from HCP participants with all four runs available were used to generate separate individual parcellations from which NSAR will be calculated. Outliers were fenced on a network basis to an upper limit of the third quartile plus 1.5 multiplied by the interquartile range, and a lower limit of the first quartile minus 1.5 multiplied by the interquartile range. An intraclass correlation coefficient was calculated comparing the NSAR from the first half of the data with the NSAR from the second half for three right-lateralized networks (Limbic-B, Visual-B, and Salience/Ventral Attention-A) and three left-lateralized networks (Language, Dorsal Attention-A, and Control-B) determined *a priori*.

#### 2.7.3 Task Effects Analysis

In the case that a large quantity of data is needed to derive a reliable estimate of lateralization, one might consider including task data in addition to any resting-state data in order to increase the amount of available data per participant. However, in this situation it would be prudent to know if task data provides the same or similar estimates as those from resting-state data. To address this concern, the NSD dataset was selected since it has a large quantity of both resting-state and task-based fMRI data per participant. Following preprocessing, a minimum of 12 resting-state runs were available for each participant, so the first 12 available resting-state runs and the first 12 available task runs were utilized (resting-state and task runs were of the same duration). Individual parcellations were then generated based on various combinations of runs within task type: even-numbered runs, odd-numbered runs, the first half of runs, the second half of runs, and two random selections of runs (without replacement). A dice coefficient was then computed to compare parcellation label overlap within task (e.g., between even and odd-numbered resting-state runs) and between tasks (e.g., between odd-numbered runs from resting-state and task runs). This comparison procedure was repeated for the NSAR intraclass correlation coefficient. Due to the non-normal nature of such a small dataset, comparisons between the task and rest parcellation dice coefficients and NSAR intraclass correlations were formally made using paired Wilcoxon Signed Rank tests (R Core Team, 2011; Wilcoxon, 1945).

### 2.8 Identifying Lateralized Networks

After establishing validity and reliability, we addressed the first hypothesis of determining whether any of the 17 networks exhibited lateralization, and of those, which were the most lateralized. The following analyses were first implemented in the HCP-Discovery dataset and then replicated in the HCP-Replication and HCPD datasets using all data available from each participant. First, to determine whether any networks exhibited lateralization, multiple regressions were implemented for each of the 17 networks. Models consisted of a given network’s NSAR value and the covariates of mean-centered age, sex, mean-centered mean framewise displacement, and handedness (measured via the Edinburgh Handedness Inventory; Oldfield, 1971). A network was considered lateralized if the model intercept was significant at the Bonferroni-corrected alpha level of 0.003. Next, to determine which networks were the most lateralized, any networks exhibiting significant lateralization in the previous tests with the same direction of lateralization were compared against each other two at a time in multiple regressions with a binary variable for the two networks and the covariates of mean-centered age, sex, mean-centered mean framewise displacement, and handedness.

### 2.9 Identifying Network Relationships

To test the second hypothesis regarding how network lateralization is potentially related between networks, a general relationship was first assessed between NSAR values averaged across like-lateralized networks followed by correlation matrices and structural equation modeling. An exploratory factor analysis (EFA) was conducted in the HCP-Discovery dataset followed by separate confirmatory factor analyses (CFAs) in the HCP-Replication, and HCPD datasets using model-adjusted lateralization values from any reliably lateralized networks. For a network to be considered reliably lateralized, it was significantly lateralized across the HCP-Discovery, HCP-Replication, and HCPD datasets. The exploratory factor analysis was chosen for its ability to identify shared relationships between the items in a data-driven manner. The *fa* function from the psych package (Revelle, 2023) was used to conduct an iterated principal factors analysis and subsequent parallel analysis. Criteria for the extraction of factors were: a minimum eigenvalue of one, visual inspection of a scree plot, and a parallel analysis. A four-factor model was hypothesized, similar to Liu et al. (2009), with each factor encompassing vision, internal thought, attention, and language. The factor structure identified in the HCP-Discovery dataset was then implemented in confirmatory factor analyses in the HCP-Replication and HCPD datasets using the *cfa* function from the lavaan package (Rosseel, 2012; Rosseel et al., 2023).

## 3 Results

### 3.1 NSAR as a Valid Measure of Lateralization

The ecological validity of NSAR was examined through comparison against laterality calculated from a language task in a subset of the HCP subjects (*N* = 221). A positive significant relationship between NSAR for the Language network and language task laterality was found (Spearman rank correlation *r* = 0.24, *p* < .001; see Figure 3).

The convergent validity of NSAR was assessed through comparison with an additional functional measure of specialization (the autonomy index) using the Spearman rank correlation. To facilitate direct comparison with NSAR values, the sign for autonomy index values was reversed. With the selected left-lateralized networks, significant relationships were found between the autonomy index and NSAR for the Language (Spearman rank correlation *r* = -0.62, *p* < .001; see Figure 3 Panel B), Dorsal Attention-A (Spearman rank correlation *r* = -0.62, *p* < .001), and the Control-B (Spearman rank correlation *r* = -0.57, *p* < .001) networks (see the top row of Figure 4). Significant relationships were also found between the autonomy index and NSAR for the selected right-lateralized networks including the Visual-B (Spearman rank correlation *r* = 0.71, *p* < .001), Salience/Ventral Attention-A (Spearman rank correlation *r* = 0.61, *p* < .001), and Limbic-B (Spearman rank correlation *r* = 0.69, *p* < .001) networks (see the second row of Figure 4). These findings indicate that NSAR and the autonomy index are measuring similar facets of specialization.

**Figure 4.**
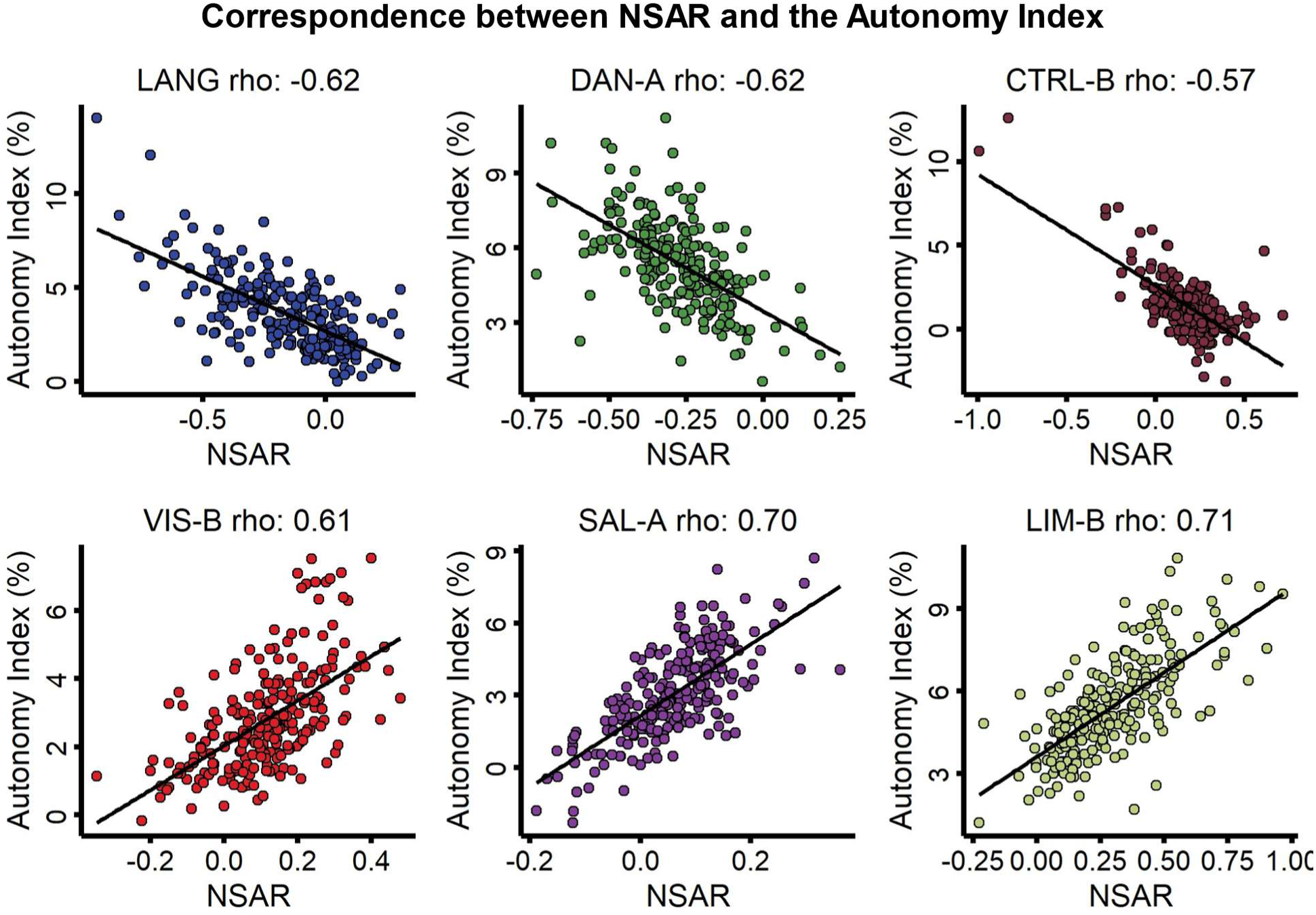
Evidence for convergent validity between the autonomy index and NSAR in a subset of HCP dataset. The top row depicts the relationships between the autonomy index and NSAR for three left-lateralized networks (Language, Dorsal Attention-A, and Control-B; Spearman rank correlation *r* = -0.57 - -0.62). The bottom row depicts the relationships between the autonomy index and NSAR for three right-lateralized networks (Visual-B, Ventral Attention-A, and Limbic-B; Spearman rank correlation *r* = 0.61 - 0.71). For each scatterplot, the line of best fit was generated using the *lm* function (no covariates) and each circle represents an individual.

Next, the external validity of NSAR was examined through comparison against two cognitive measures using a CCA: a reading task (ORRT) and an attention/inhibitory control task (the Flanker task). In preparation for the CCA in a subset of HCP participants (*N* = 232; no missing data), linearity and heteroskedasticity of age-, sex-, handedness-, and mean framewise displacement-adjusted NSAR values from eight significantly lateralized networks and the age-and sex-adjusted values from two cognitive measures were evaluated in pairwise plots, which were followed by the Doornik-Hansen multivariate test for normality (*DH.test* function from the mvnTest package; *DH* = 164.21, *p* = 0; Doornik & Hansen, 2008; Pya et al., 2016). Tests of dimensionality for the CCA indicated that one of the two canonical dimensions was statistically significant at the .05 level. This dimension had a canonical correlation of 0.34 (*F*(16, 444) = 0.87, *p* = .008) between the cognitive measures and NSAR values, while the canonical correlation was much lower for the second, nonsignificant dimension at 0.14 (*F*(7, 223) = 0.98, *p* = .75). Table 1 presents the standardized canonical coefficients for the first dimension across the cognitive measures and eight lateralized networks. Of the cognitive variables, the first canonical dimension was most strongly influenced by language ability (β (standardized canonical coefficient) = -0.99). In terms of lateralized networks, the Visual-B (β = -0.33, *r* = -0.13), Language (β = 0.39, *r* = 0.2), Dorsal Attention-A (β = -0.54, *r* = -0.17), and Control-C (β = 0.48, *r* = 0.13) networks appeared to contribute the most to the first canonical dimension. Haufe-transformed feature weights indicated that for every one-unit increase in Language network lateralization, the first dimension, representing language ability, increases by 0.13 (see Table 1). These findings suggest that there is a relationship between network lateralization and cognitive abilities, specifically language.

**Table 1.** Canonical Correlation Analysis Results for Dimension 1in a Subset of the HCP Dataset (N = 232)

|  | Standardized<br>Canonical<br>Coefficient | Haufe-Transformed<br>Weight | Correlation with<br>Canonical Variate* |
| --- | --- | --- | --- |
| Cognitive Variables |  |  |  |
| Language (ORRT) | -0.99 | -0.45 | **-0.32** |
| Attention (Flanker) | 0.39 | 2.22 | 0.07 |
| Lateralized Networks |  |  |  |
| Visual-B | -0.33 | -0.05 | **-0.13** |
| Language | 0.39 | 0.13 | **0.2** |
| Dorsal Attention-A | -0.54 | -0.07 | **-0.17** |
| Salience/Ventral<br>Attention-A | -0.18 | -0.03 | -0.1 |
| Control-B | 0.11 | -0.01 | -0.02 |
| Control-C | 0.48 | 0.04 | **0.13** |
| Default-C | 0.37 | 0.05 | 0.12 |
| Limbic-B | -0.17 | -0.03 | -0.05 |

\*Bolted values were significant at the $p < .05$ level following multiple comparison corrections.

### 3.2 NSAR as a Reliable Measure of Lateralization

#### 3.2.1 Stable Estimate Analysis

To address the question of how much data is needed in order to obtain a stable estimate of NSAR values, combinations of five-minute increments (5, 10, 15…30 minutes) were compared against 30 independent minutes of data in a subset of HCP subjects. The intraclass correlations indicate that only five minutes of data are needed to obtain moderate to good intraclass correlations for the majority of subjects (see Figure 5 Panel A). Of note, poor and excellent interclass correlations were observed for some subjects. The stable estimate analysis was also approached from a network basis (as opposed to the subject basis presented in Figure 5 Panel A). Networks with the lowest intraclass correlations included the Limbic-A and Control-A networks, while networks with the greatest intraclass correlations included Visual-A, Limbic-B, and Default-A (for overall distributions, see Figure 5 Panel B; for specific network intraclass correlation coefficients, see Supplementary Figure S2). Interestingly, not all networks improved in reliability with additional data, including the Limbic-A and Control-A networks. This is likely a reflection of a poor signal-to-noise ratio. For parcellation label overlap estimates, see Supplementary Figure S3.

**Figure 5.**
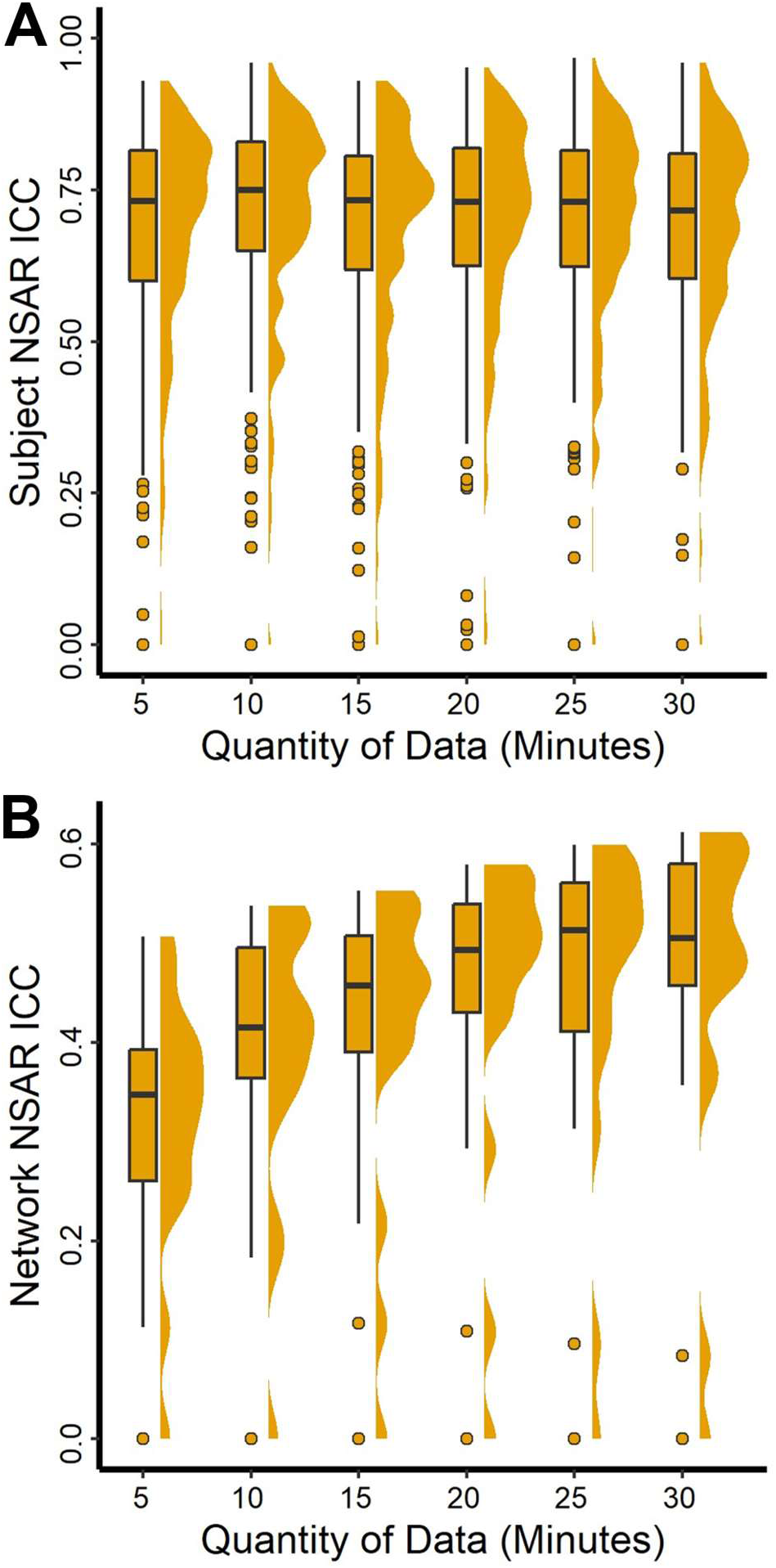
Evidence for reliable estimates of NSAR in the HCP dataset. Panel A depicts the intraclass correlation coefficient calculated for each subject’s 17 NSAR values for each time increment (5, 10, 15 … 30 minutes) and the subject’s 17 NSAR values from 30 independent minutes of data. Panel B depicts the intraclass correlation coefficient calculated for each network’s mean NSAR value between the 30 independent minutes of data and each increment of data. The distribution of intraclass correlation coefficients is shown for the 17 networks. Specific network intraclass correlation coefficients are displayed in Supplementary Figure S2.

#### 3.2.2 Test-Retest Reliability Analysis

Using HCP subjects with all four resting-state runs available post-preprocessing (*N* = 232), test-retest reliability was assessed for three left-lateralized networks (Language, Dorsal Attention-A, and Control-B) and three right-lateralized networks (Limbic-B, Visual-B, and Salience/Ventral Attention-A) determined *a priori*. For the left-lateralized networks, intraclass correlations were within the moderate range, from 0.56 to 0.63, with the lowest being the Dorsal Attention-A network (ICC = 0.56, *F*(231, 231) = 3.6, *p* < .001, 95% CI [0.47, 0.64]; see Figure 6). For the right-lateralized networks, intraclass correlations remained in the moderate range, between 0.58 to 0.71, with the Visual-B network exhibiting the lowest reliability (ICC = 0.58, *F*(231, 231) = 3.7, *p* < .001, 95% CI [0.48, 0.66]).

**Figure 6.**
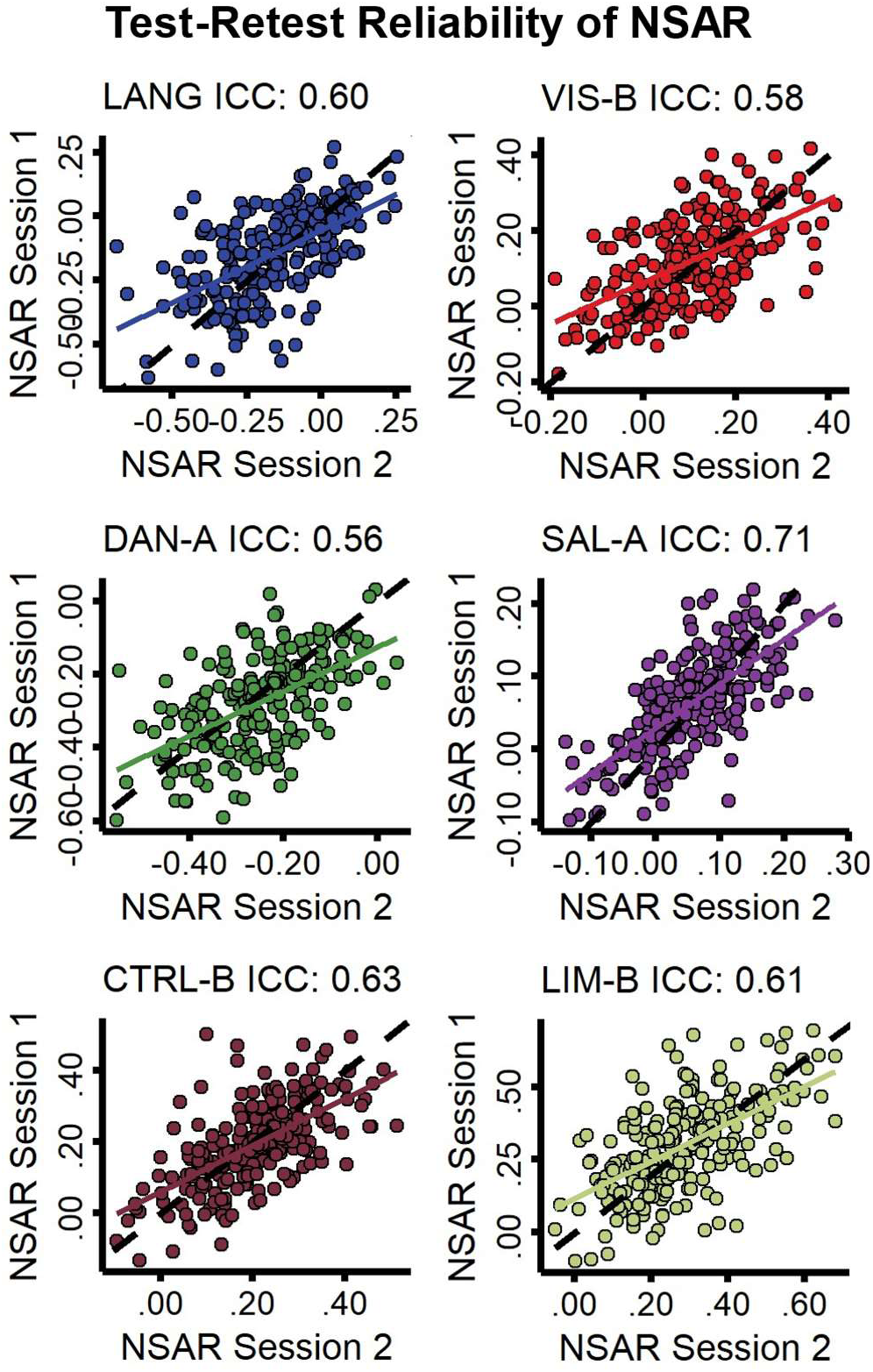
Test-retest reliability of NSAR values for left-and right-lateralized networks in 232 HCP subjects. Left-lateralized networks (left column) included Language, Dorsal Attention-A, and Control-B. Right-lateralized networks (right column) included Visual-B, Salience/Ventral Attention-A, and Limbic-B. In each plot, a circle represents a subject.

#### 3.2.3 Task Effects on Individual Parcellations and NSAR

Using the NSD dataset (*N* = 8) to compare potential differences between resting-state and task fMRI on individual parcellations and NSAR estimates, we found differences between the within-task comparisons and between task comparisons for both the parcellation dice coefficients and NSAR intraclass correlations (see Figure 7). Wilcoxon signed rank comparisons revealed a difference in within-task (Task-Task and Rest-Rest) dice coefficients for even versus odd numbered runs (*V* = 36, *p* = .008), but no difference for the first half versus the second half of runs (*V* = 29, *p* = .15) or the random selection of runs (*V* = 31, *p* = .08). Regardless of how the data were split, a task effect in dice coefficient was found between within-task (Task-Task) and between-task (Task-Rest) dice coefficients for even versus odd numbered runs (*V* = 36, *p* = .008), the first half versus the second half of runs (*V* = 36, *p* = .008), and the random selection of runs (*V* = 36, *p* = .008).

**Figure 7.**
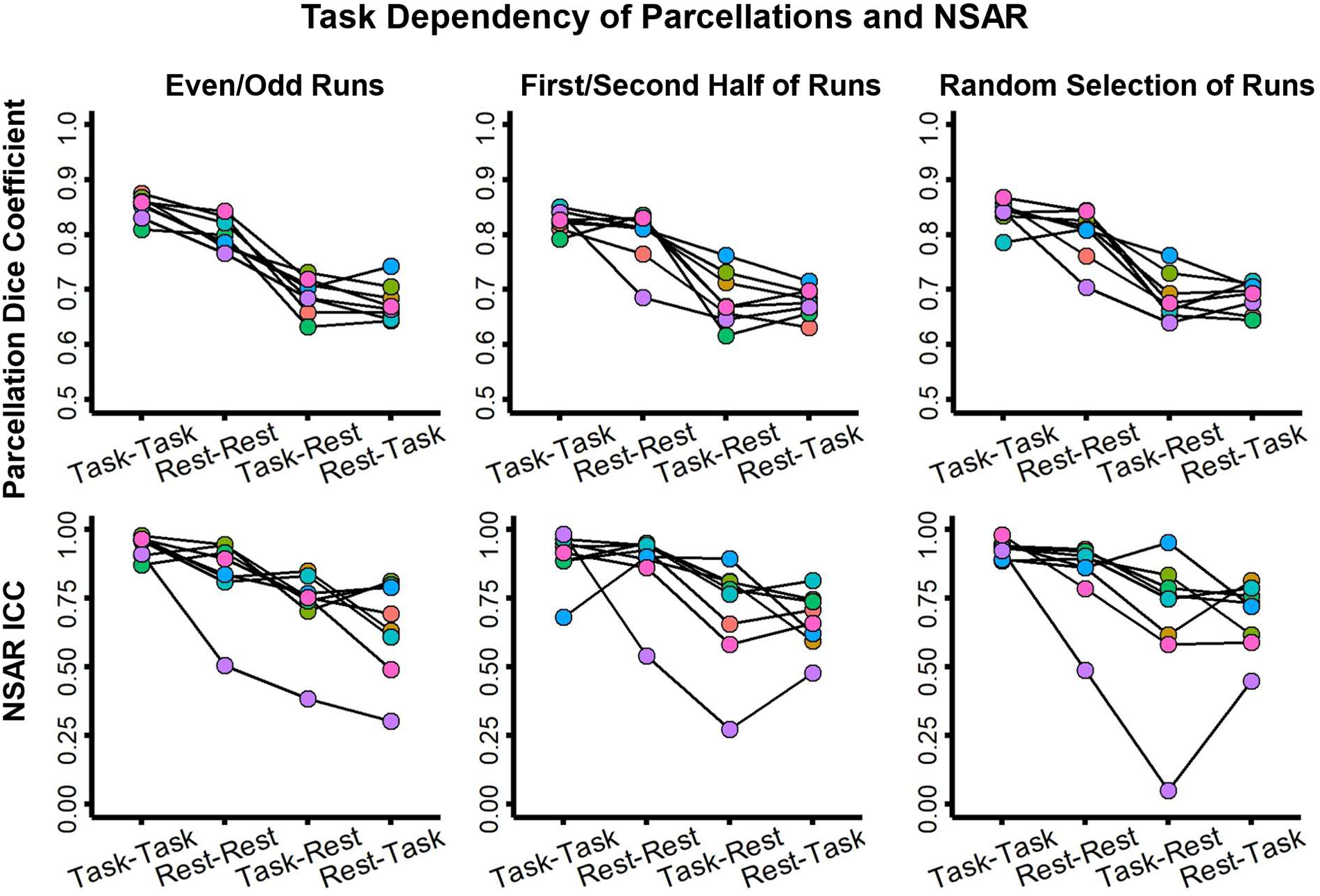
Task dependency of individual parcellations and NSAR in the NSD dataset. Depicted in the top row are the dice coefficients for the individual parcellations between 30-minute increments of resting-state or task fMRI data. Regardless of how the data were split (even-versus odd-numbered runs, the first half versus the second half, or a random selection without replacement), a task effect was found. Depicted in the second row are the NSAR intraclass correlation coefficients computed in individuals across networks. In each plot, circles connected by a line represent an individual.

Similarly, with the NSAR intraclass coefficients, no significant difference was found for within-task (Task-Task and Rest-Rest) reliability across the even versus odd numbered runs (*V* = 31, *p* = .08) and the first half versus the second half of runs (*V* = 19, *p* = .95), but not for the random selection of runs (*V* = 35, *p* = .02). However, a significant difference was not found between within-task (Task-Task) and between-task (Task-Rest) intraclass correlation coefficients across the even versus odd numbered runs (*V* = 31, *p* = .08), the first half versus the second half of runs (*V* = 31, *p* = .08), but for the random selection of runs (*V* = 34, *p* = .02).

### 3.3 Networks with the Greatest Lateralization

To test the first hypothesis that networks associated with language, visuospatial attention, and executive control would show the greatest hemispheric lateralization, networks were first evaluated for lateralization and then compared against each other. To begin, a series of multiple regressions were used to identify if any of the 17 networks were lateralized, first in the HCP-Discovery dataset and then in the HCP-Replication and HCPD datasets. Networks with significant lateralization (*p* < .003) in the same direction (e.g., right or left lateralization) across all three datasets included nine networks, of which four were left-lateralized (Language, Dorsal Attention-A, Control-A and Default-C) and five were right-lateralized (Visual-B, Salience/Ventral Attention-A, Control-B, Control-C, and Limbic-B; see Supplementary Table 1).However, given the very low reliability of the left-lateralized Control-A network (mean ICC = 0.12; see Supplementary Figure S2), this network was not considered further. None of the covariates were reliably significant for a given network across all three datasets. See Figure 8 for model-adjusted NSAR values for each of the 17 networks and see Figure 9 for the percentage of surface area occupied by the eight most lateralized networks.

**Figure 8.**
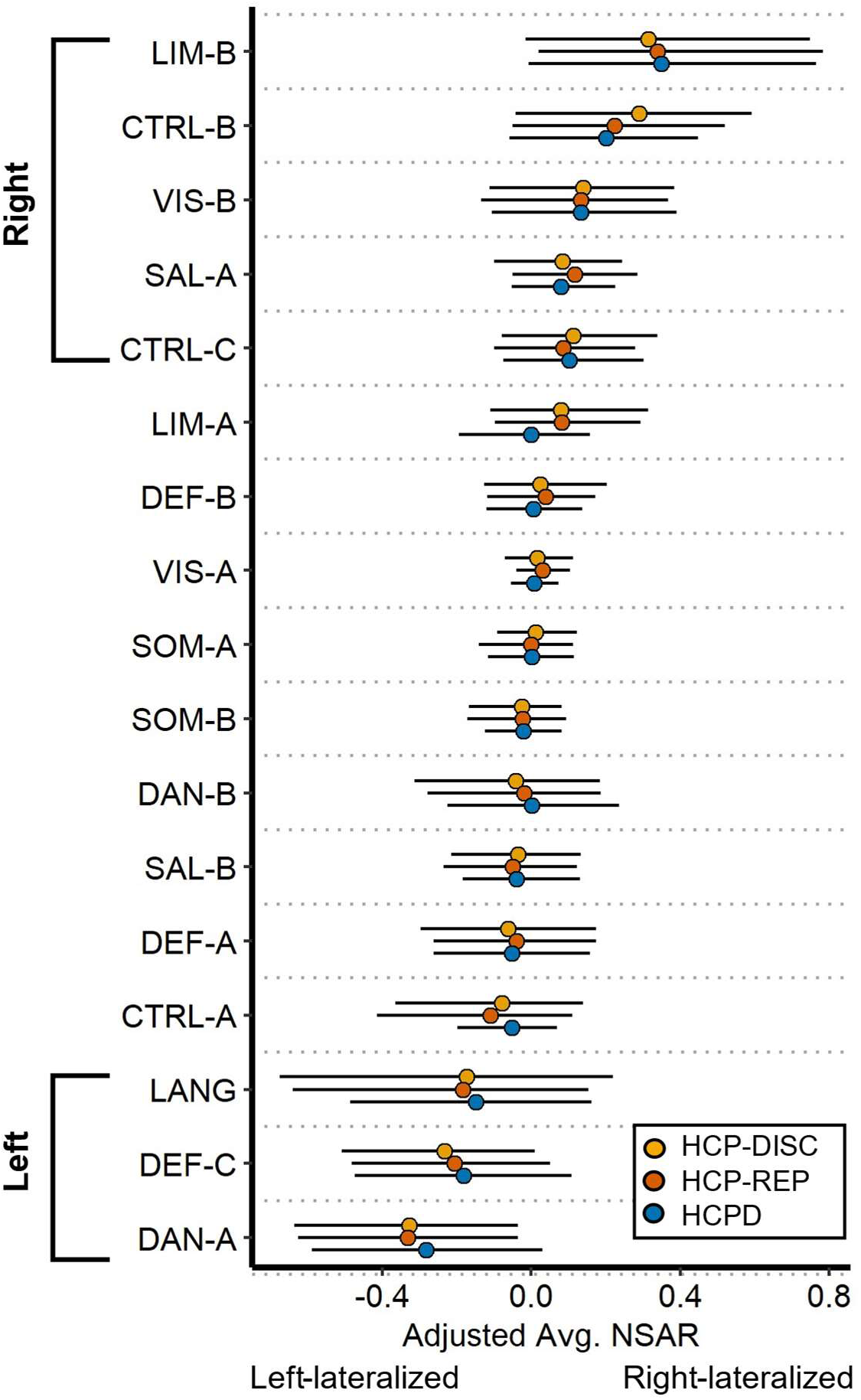
Lateralization for 17 networks across the HCP-Discovery, HCP-Replication, and HCPD datasets. On the y-axis are the 17 networks and on the x-axis are the adjusted NSAR values, with negative values representing left hemisphere lateralization and positive values representing right hemisphere lateralization. Bars represent the 2.5 and 97.5 percentiles. NSAR values were adjusted by regressing out the effects of mean-centered age, mean-centered mean framewise displacement, and sex using the following formula: NSAR_adjusted_ = NSAR_raw_ — [β_1_(mean-centered age_raw_ – mean of mean-centered age_raw_) + β_2_(mean-centered FD_raw_ – mean of mean-centered FD_raw_) + β_3_(sex_raw_ – mean sex_raw_) + β_4_(handedness_raw_ – mean handedness_raw_)]. NSAR adjustment occurred separately for each network within each dataset. Lines represent the standard error. Across the three datasets, eight networks were reliably and significantly lateralized (left-lateralized: Language, Dorsal Attention-A, and Default-C; right-lateralized: Visual-B, Salience/Ventral Attention-A, Control-B, Control-C, and Limbic-B).

**Figure 9.**
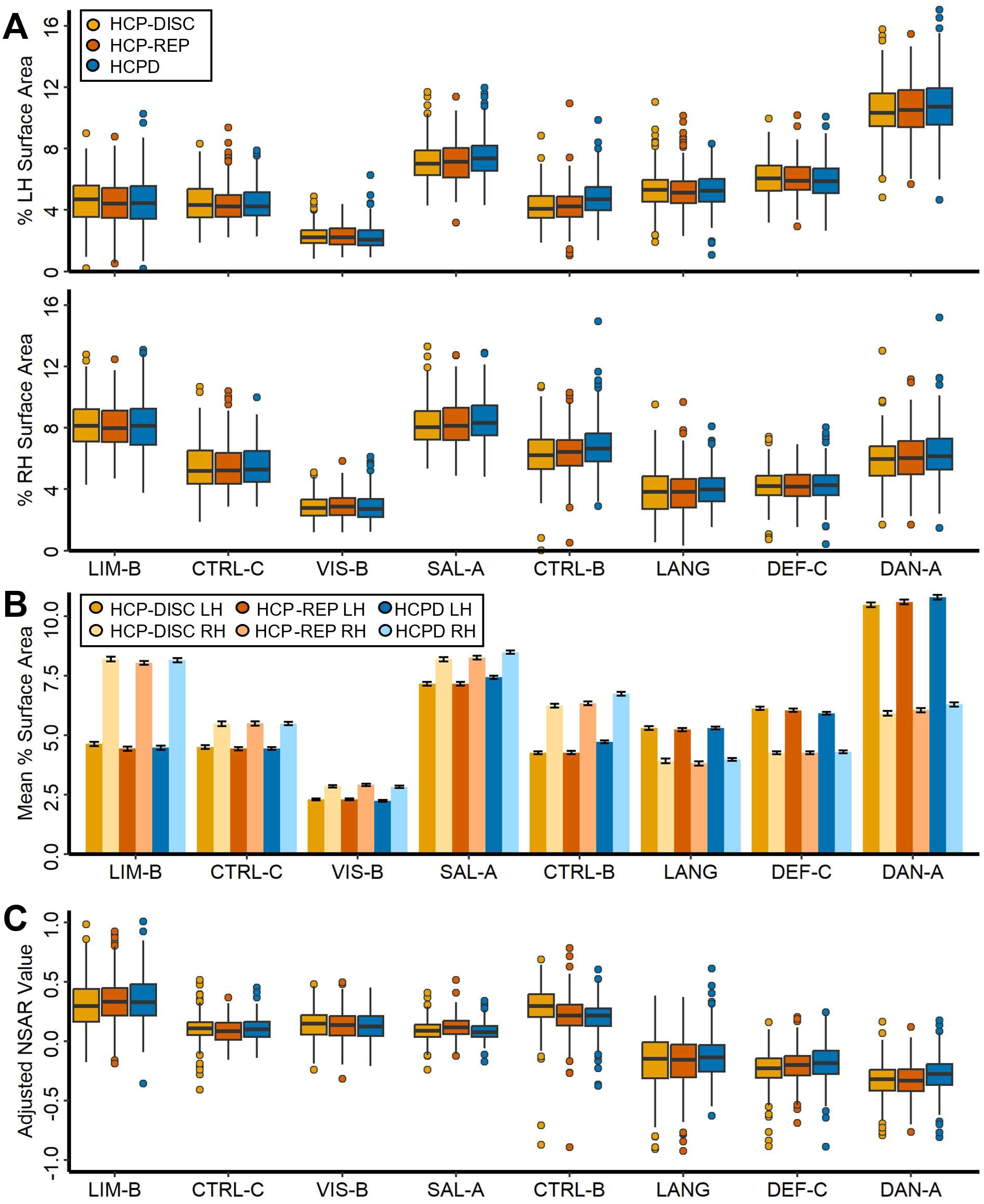
Percent surface area for 8 lateralized networks across the HCP-Discovery, HCP-Replication, and HCPD datasets. Depicted in the top of Panel A is the percentage of the surface area occupied by a given lateralized network for the left hemisphere (top panel) and right hemisphere (bottom panel). Depicted in Panel B is the mean percentage of surface area occupied by a lateralized network, with standard error bars. The left and right hemisphere estimates are displayed side-by-side for each dataset. In Panel C, the adjusted NSAR values for each network are shown. In Panels A and C, points represent individual outliers.

Following the identification of eight lateralized networks, a series of multiple regressions were used to compare networks with the same direction of lateralization two at a time in order to identify the networks with the greatest lateralization. Models included a binary network variable and the covariates of mean-centered age, sex, handedness, and mean-centered mean framewise displacement. Of the left-lateralized networks, the Dorsal Attention-A network was the most lateralized compared with the Language and Default-C networks, and this pattern was replicated across the HCP-Discovery, HCP-Replication, and HCPD datasets (see Supplementary Table 2). Of the right-lateralized networks, the Limbic-B network was the most lateralized, followed by the Control-B network, Visual-B and Control-C networks (not significantly different), and the Salience/Ventral Attention-A network. This pattern was replicated across the three datasets as well (see Supplementary Table 3). Contrary to our hypothesis that networks associated with language, visuospatial attention, and executive control would show the greatest lateralization, we identified the Dorsal Attention-A network as the most left-lateralized and the Limbic-B network as the most right-lateralized.

### 3.4 Relationships between Networks’ Lateralization

Next, we investigated how lateralization in one network may influence the lateralization of other networks. This second hypothesis was assessed first through general correlations followed by both correlation matrices and structural equation modeling conducted in triplicate across the HCP-Discovery, HCP-Replication, and HCPD datasets. First, model-adjusted NSAR values were averaged across like-lateralized networks before the averaged left-lateralized values (from the Language, Dorsal Attention-A, and Default-C networks) were correlated with the averaged right-lateralized values (from the Visual-B, Salience-Ventral Attention-A, Control-B, Control-C, and Limbic-B networks). A general negative relationship between left-lateralized and right-lateralized networks was found across each dataset (HCP-Discovery: *r*(274) = -0.67, *p* < .001; HCP-Replication: *r*(275) = -0.59, *p* < .001; HCPD: *r*(343) = -0.66, *p* < .001). Next, correlation matrices of the model-adjusted NSAR values from the eight lateralized networks evidenced moderate negative relationships between the left-and right-lateralized networks across individuals (see Figure 10). In the HCP-Discovery dataset, negative relationships were found between the Limbic-B and Dorsal Attention-A networks (*r*(274) = -0.45, *p* < .001, 95% CI [-0.54, -0.36]; see Figure 11 Panel A), the Limbic-B and Default-C networks (*r*(274) = -0.42, *p* < .001 , 95% CI [-0.51, -0.31]; see Figure 11 Panel B), the Default-C and Visual-B networks (*r*(274) = -0.16, *p* = .007, 95% CI [-0.28, -0.05]), the Default-C and Control-B networks (*r*(274) = -0.27, *p* < . 001, 95% CI [-0.38, -0.16]), the Default-C and Control-C networks (*r*(274) = - 0.17, *p* = .004, 95% CI [-0.29, -0.06]), the Control-B and Language networks (*r*(274) = -0.31, *p* < .001, 95% CI [-0.41, -0.19]), and the Language and Salience/Ventral Attention-A networks (*r*(274) = -0.3, *p* < .001, 95% CI [-0.41, -0.19]; see Figure 11 Panel C). Interestingly, a negative relationship was also found between two left-lateralized networks: Dorsal Attention-A and Language (*r*(274) = -0.23, *p* < .001, 95% CI [-0.34, -0.11]). Each negative relationship was replicated across the HCP-Replication and HCPD datasets (see Figure 10). These relationships support the dependent hypothesis, which suggests that having one highly lateralized network corresponds with increased lateralization in other networks within the individual, and that this pattern is systematic across individuals.

**Figure 10.**
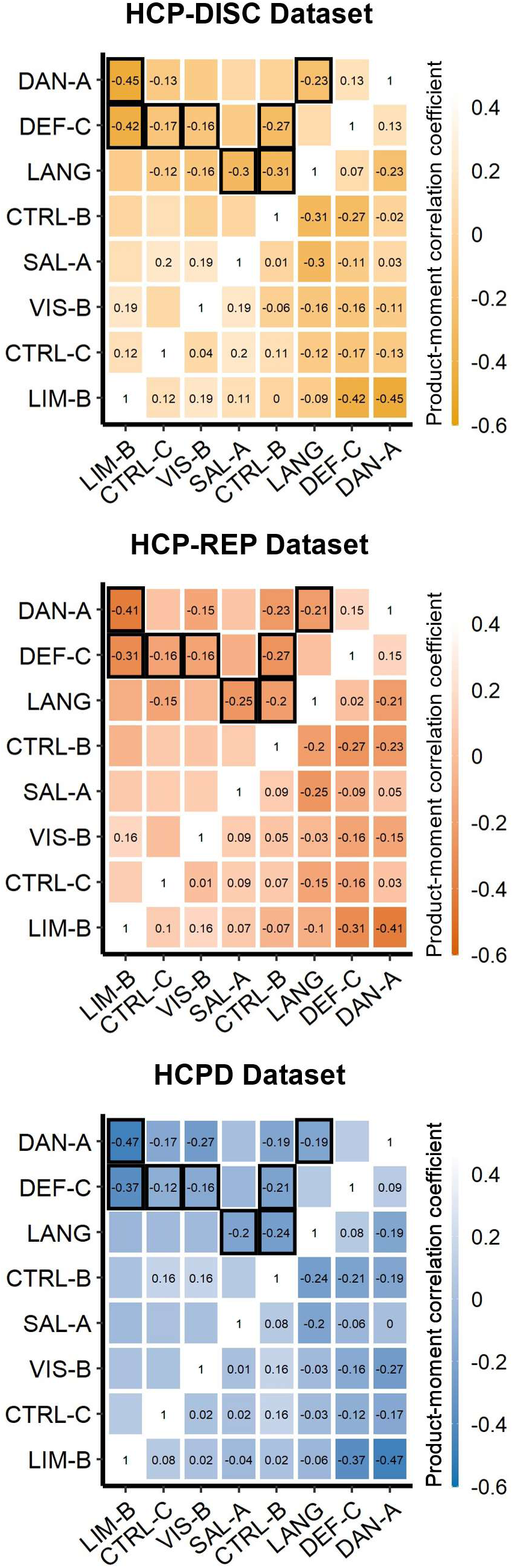
Relationships between lateralized networks across the HCP-Discovery, HCP-Replication, and HCPD datasets. Correlation matrices were created from the model-adjusted NSAR values from the eight lateralized networks (Visual-B, Language, Dorsal Attention-A, Salience/Ventral Attention-A, Control-B, Control-C, Default-C, and Limbic-B), controlling for sex, mean-centered age, mean-centered framewise displacement, and handedness. Correlation values thresholded at *p* = .05 are displayed in the upper triangles, and consistent relationships have been highlighted with black boxes.

**Figure 11.**
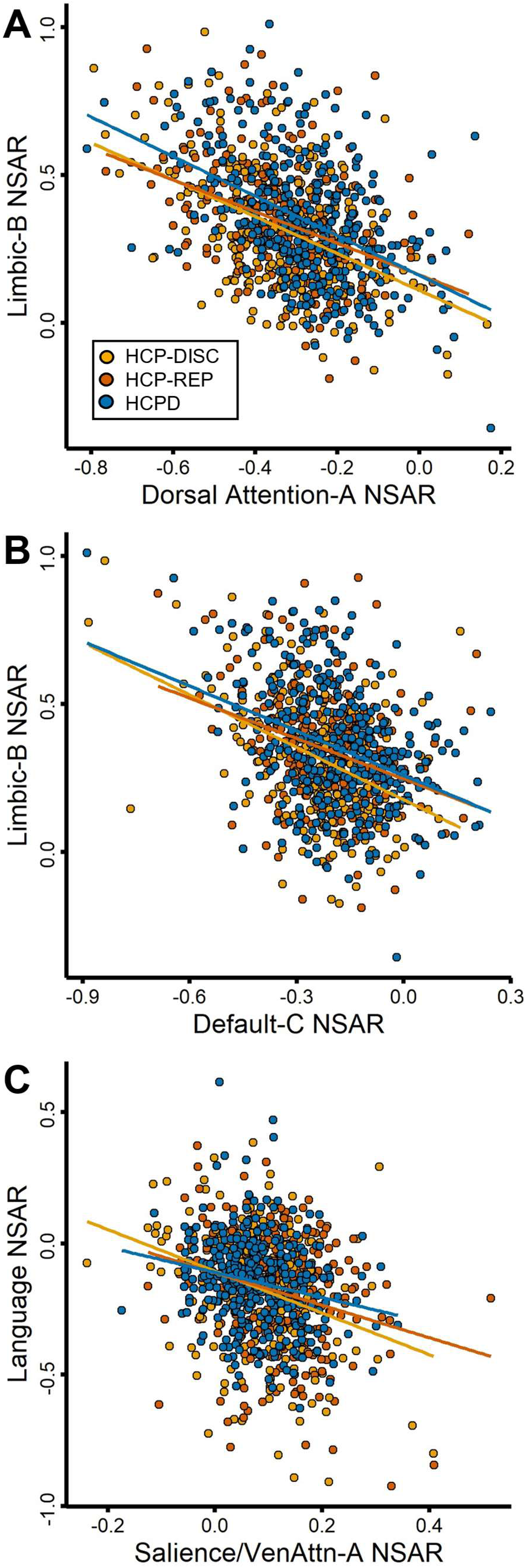
Negative correlations between highly left-and right-lateralized networks across the HCP-Discovery, HCP-Replication, and HCPD datasets. Panel A depicts the negative relationship between the Limbic-B and Dorsal Attention-A networks (HCP-Discovery: *r*(274) = -0.45, adjusted *R^2^* = 0.2; HCP-Replication: *r*(275) = -0.41, adjusted *R^2^* = 0.16; HCPD: *r*(341) = -0.47, adjusted *R^2^* = 0.22). Panel B depicts the negative relationship between the right-lateralized Limbic-B and left-lateralized Default-C networks (HCP-Discovery: *r*(274) = -0.42, adjusted *R^2^* = 0.17; HCP-Replication: *r*(275) = -0.31, adjusted *R^2^* = 0.09; HCPD: *r*(341) = -0.37, adjusted *R^2^* = 0.14). Panel C depicts the negative relationship between the right-lateralized Salience/Ventral Attention-A network and left-lateralized Language network (HCP-Discovery: *r*(274) = -0.3, adjusted *R^2^* = 0.09; HCP-Replication: *r*(275) = -0.25, adjusted *R^2^* = 0.06; HCPD: *r*(341) = -0.2, adjusted *R^2^* = 0.04). In each panel, a circle represents a single participant’s model-adjusted NSAR value, which was adjusted for mean-centered age, sex, handedness, and mean-centered mean framewise displacement.

#### 3.4.1 EFA in the HCP-Discovery Dataset

As an additional method for exploring relationships between lateralized networks, an EFA was implemented in the HCP-Discovery dataset, followed by CFAs in the HCP-Replication, and HCPD datasets. In preparation for the EFA in the HCP-Discovery dataset (*N* = 276; no missing data), linearity and heteroskedasticity of adjusted NSAR values from the eight lateralized networks were evaluated in pairwise plots, which were followed by the Doornik-Hansen multivariate test for normality (*DH.test* function from the mvnTest package; *DH* = 202.89, *p* = 0; Doornik & Hansen, 2008; Pya et al., 2016). The NSAR values were then evaluated for multicollinearity, and no items had Variance Inflation Factor values greater than 1.65 (*vif* function from the psych package; Revelle, 2023). Additional assumptions testing included Bartlett’s test of sphericity and the Kaiser-Meyer-Olkin (KMO) Measure of Sampling Adequacy. For the test of sphericity, we rejected the null hypothesis that there is no correlation among the items (χ^2^(28) = 293.43, *p* < .001). Additionally, the KMO test was .46, revealing that the extracted factors will account for an unacceptable amount of common variance.

To examine network relationships, a principal factors analysis in the HCP-Discovery dataset was performed. Using the correlation matrix from eight lateralized networks, two factors were extracted. This first factor had an eigenvalue of 1.38 (explaining 57% of the variance; see Table 2 for factor loadings) and the second factor had an eigenvalue of 1.02 (explaining 43% of the variance). Of note, the left-lateralized networks load negatively onto the first extracted factor while right-lateralized networks load positively, suggesting that this factor encompasses right-hemisphere lateralization, with the opposite in the second extracted factor.

**Table 2.** Summary of Exploratory Factor Analysis Results for the NSAR Scores Using Iterated Principal Factors in the HCP-Discovery Dataset (N = 276)

| Network | Factor 1 Loadings | Factor 2 Loadings |
| --- | --- | --- |
| Limbic-B | **0.73** | -0.32 |
| Control-C | 0.28 | 0.06 |
| Visual-B | 0.29 | 0.04 |
| Salience/VenAttn-A | 0.28 | 0.26 |
| Control-B | 0.22 | 0.23 |
| Language | -0.38 | **-0.75** |
| Default-C | **-0.51** | 0.05 |
| Dorsal Attention-A | -0.37 | **0.48** |
| Eigenvalues | 1.38 | 1.02 |
| Proportion of variance explained | 0.57 | 0.43 |

*Note:* Factor loadings over .40 appear in bold.

#### 3.4.2 CFAs in the HCP-Replication and HCPD Datasets

In preparation for the CFA in the HCP-Replication dataset (*N* = 277; no missing data), linearity and heteroskedasticity of adjusted NSAR values were evaluated in pairwise plots, which were followed by the Doornik-Hansen multivariate test for normality (*DH* = 43.29, *p* < .001; Doornik & Hansen, 2008; Pya et al., 2016). The NSAR values were then evaluated for multicollinearity, and no items had Variance Inflation Factor values greater than 1.3. Additional assumptions testing included Bartlett’s test of sphericity and the Kaiser-Meyer-Olkin (KMO) Measure of Sampling Adequacy. For the test of sphericity, we rejected the null hypothesis that there is no correlation among the items (χ^2^(6) = 101.53, *p* < .001). Additionally, the KMO test was .49, revealing that the extracted factors will account for an unacceptable amount of common variance. This process of evaluating assumptions was also performed in the HCPD dataset (*N* = 343; no missing data), starting pairwise plots and the Doornik-Hansen multivariate test for normality (*DH* = 44.37, *p* < .001). Multicollinearity was then evaluated, and no items had Variance Inflation Factor values greater than 1.49. Additionally, for Bartlett’s test of sphericity, we rejected the null hypothesis that there is no correlation among the items (χ^2^(6) = 164.59, *p* < .001). Furthermore, the KMO test was 0.47, revealing that the extracted factors will account for an unacceptable amount of common variance.

To examine network relationships and potentially replicate the HCP-Discovery EFA, a confirmatory factor analyses were performed in the HCP-Replication and HCPD datasets using the *cfa* function from the lavaan package (Rosseel, 2012; Rosseel et al., 2023). The structural model consisted of two factors, with Limbic-B and Default-C loaded onto the first factor and Language and Dorsal Attention-A loaded onto the second factor. In the HCP-Replication dataset, the model provided fair fit to the data: χ^2^(2) = 61.95, *p* < .001; confirmatory fit index (CFI) = 0.38; root-mean-square error of approximation (RMSEA) = 0.33; standardized root mean square residual (SRMR) = 0.14. Similar results were found in the HCPD dataset, for which the model provided fair fit to the data: χ^2^(2) = 102.02, *p* < .001; CFI = 0.38; RMSEA = 0.38; SRMR = 0.16. Standardized loadings for each network across both CFAs are shown in Table 3.

**Table 3.**
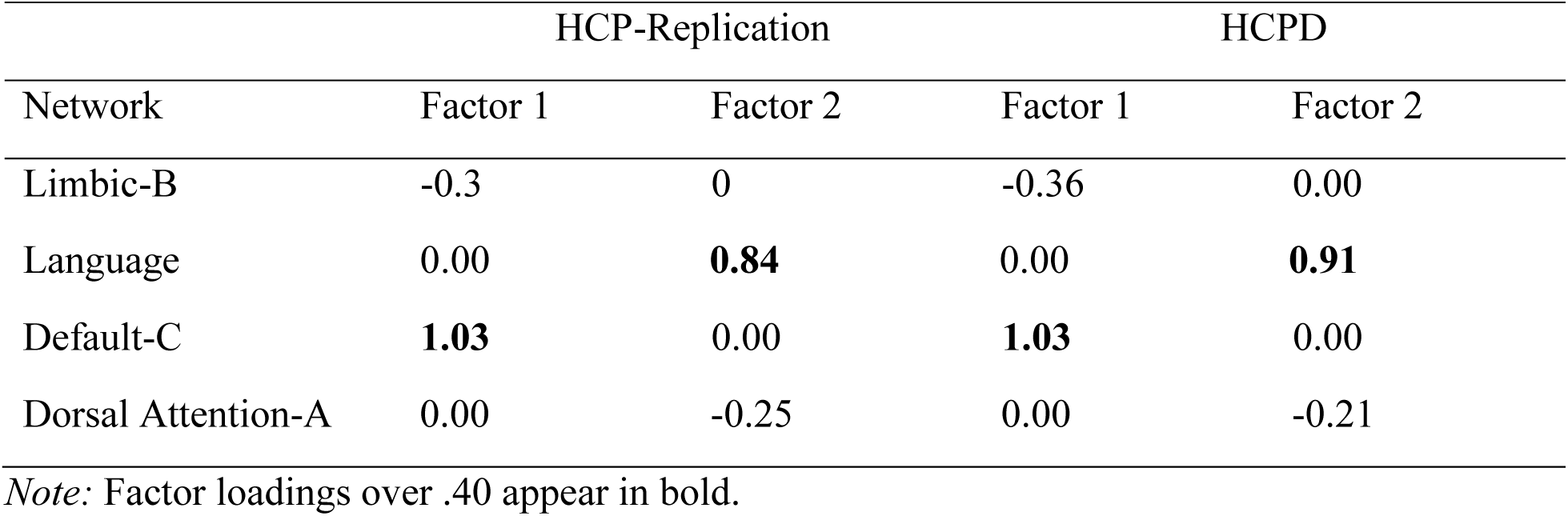
Standardized Loadings for a Two-Factor Confirmatory Factor Analysis Model of NSAR Scores in the HCP-Replication (N = 277) and HCPD (N = 343) Datasets.

## 4 Discussion

In this study, we implemented a novel measure of lateralization based on high-resolution individual network parcellations (NSAR). Using NSAR, we identified eight networks that were reliably lateralized across three independent datasets. Furthermore, we examined potential relationships between networks’ NSAR values and found evidence supporting a dependent hypothesis of lateralization. These findings shed new light on hemispheric specialization, which has implications for the understanding of brain organization and development (Toga & Thompson, 2003), individual differences (Perez et al., 2023), human-defining cognitive processes (Hartwigsen et al., 2021), and neurodevelopmental conditions (Eyler et al., 2012; Kong et al., 2022). Previously, functional lateralization has been assessed through a variety of approaches dependent on intrinsic connectivity, primarily at the group level. However, recent evidence suggests that group-based approaches can obfuscate the idiosyncratic anatomy of individuals and blur potentially meaningful and clinically useful variability (Gratton et al., 2020; Lynch et al., 2020; Salvo et al., 2021). For example, the language network has high spatial variability across individuals (Braga et al., 2020; Fedorenko, Duncan, et al., 2012), holding ramifications for the accurate assessment of lateralization for this and other variable networks.

### 4.1 Evidence for the Validity and Reliability of NSAR

In this study, we examined functional lateralization using a novel surface area-based index. This measure was developed methodologically through the examination of ecological, convergent, and external validity, as well as a stable estimate analysis, test-retest reliability, and potential task effects. Notably, language task laterality appears to have a positive relationship with Language network NSAR, suggesting that there is a degree of concordance between this resting-state measure of laterality and a task-based measure of laterality. Furthermore, estimates from this surface area approach to lateralization appear to converge with a different functional connectivity-based method (the autonomy index). This result supports the idea that NSAR is capturing lateralization in a way that is valid while being distinct from the autonomy index in how it is derived. Unlike the autonomy index, the formula for NSAR does not normalize for brain size or deal in the minutiae of individual functional connections. Rather, NSAR is calculated based on a network’s surface area. Additionally, potential relationships between network NSAR values and two cognitive measures were investigated in an analysis of external validity. Interestingly, a relationship between the laterality of the Visual-B, Language, Dorsal Attention-A, Control-C, and Default-C networks and language ability was identified. Similarly, others have identified a link between language function and left hemisphere lateralization during language production (Groen et al., 2012), and between the lateralization of functional networks and their associated cognitive abilities (Gotts et al., 2013).

Reliability analyses indicated that NSAR is stable within individuals, even after just five minutes of resting-state fMRI data. Interestingly, networks with the greatest reliabilities included the visual and somatomotor networks. This is in keeping with Kong et al. (2019), who found that sensorimotor networks exhibited lower inter-subject functional connectivity variability than association networks. Since NSAR is indirectly based on an individual’s functional connectivity profiles, this result is unexpected.

In addition to the quantity of data available per participant, we also examined the effect of data type (task versus rest) on NSAR estimates within individuals. While within-task type reliability was high, we found that there was indeed a task effect such that resting-state fMRI and task fMRI did not yield identical parcellations and NSAR estimates within individuals. This finding supports the hypothesis that resting-state fMRI can be thought of as another arbitrary task state (Buckner et al., 2013). Yet, the “task” of resting-state fMRI can result in greater variability in functional connectivity compared with that resulting from task fMRI, perhaps resulting from mind wandering (Elton & Gao, 2015). And when predicting individual traits, task-based models outperform rest-based models, with this difference likely reflecting the “unconstrained nature” of the resting state (Greene et al., 2018). Since NSAR estimates are derived from individual parcellations which are in turn generated from individual functional connectivity profiles, it stands to reason that connectivity differences resulting from task type could trickle down to differences in NSAR estimates.

### 4.2 The Identification of Eight Reliably Lateralized Networks

Following the methodological development of NSAR, we reliably identified eight lateralized networks across three datasets: Visual-B, Language, Dorsal Attention-A, Salience/Ventral Attention-A, Control-B, Control-C, Default-C, and Limbic-B. While a ninth lateralized network was reliably identified (Control-A), this network was discarded from further analysis due to very poor reliability. Previously, several of these networks have been established as lateralized, particularly those associated with language and visuospatial attention processing.

#### 4.2.1 The Dorsal Attention-A Network Exhibited the Greatest Left-Lateralization

Previously, left-lateralized networks have included the language, frontoparietal control, and default networks. More specifically, evidence for the lateralization of the language network has been derived from a variety of methods including the Wada test (Desmond et al., 1995; Wada & Rasmussen, 1960), lesion cases (Broca, 1861; Wernicke, 1995), task fMRI (Elin et al., 2022; Fedorenko, Duncan, et al., 2012; Fedorenko et al., 2010, 2011; Fedorenko, McDermott, et al., 2012; Lipkin et al., 2022; Malik-Moraleda et al., 2022; Olulade et al., 2020; Scott et al., 2017; Wilson et al., 2017), and resting-state fMRI (Braga et al., 2020; Labache et al., 2020; Zhu et al., 2014), among others. Using NSAR, we also identified the language network as being strongly left-lateralized. However, unlike a prior comparative study (Braga et al., 2020), which examined lateralization in the language, salience, default, and frontoparietal networks (but not a dorsal attention network), we did not find that the language network was the most left-lateralized network. Instead, we identified the Dorsal Attention-A network as being the most left-lateralized. Unlike the ventral attention network, the dorsal attention network has been previously identified as a bilateral network (Fox et al., 2006; for review see Mengotti et al., 2020). This was the case for the Dorsal Attention-B network, which was not a significantly lateralized network across the three datasets. However, there is evidence for a left-lateralized dorsal attention network across both left-and right-handed individuals, stemming from a within-individual network variants approach (see Figure 7 Panel C of Perez et al., 2023). Additionally, it could be that a finer-grained parcellation deconstructs the dorsal attention network into one bilateral and one lateralized network, similar to previous within-individual work on the default network (Braga & Buckner, 2017; DiNicola et al., 2020).

#### 4.2.2 Replication of Right-Lateralized Attention, Control, and Limbic Networks

This is not the first study to identify the ventral attention, control, and limbic networks as being lateralized. Abundant evidence exists for the right-lateralization of visuospatial/ventral attention, stemming from task fMRI (Beume et al., 2015; Cai et al., 2013; Jansen et al., 2004; Shulman et al., 2010; Siman-Tov et al., 2007; Umarova et al., 2010; J. Wang et al., 2016; Zago et al., 2016, 2017), resting-state fMRI (Braga et al., 2020; Wang et al., 2014), hemispatial neglect cases (Corbetta & Shulman, 2011), and others (for review, see Mengotti et al., 2020). Interestingly, we identified the Salience/Ventral Attention-A but not the Salience/Ventral Attention-B network as being right-lateralized. Once more, this may be due to the network resolution selected (*k* = 17), which may have split the canonical ventral attention network into a bilateral and a right-lateralized network.

While this study successfully replicated right-lateralized control networks (Control-B and Control-C), a left-lateralized control network was not identified. Previously, Wang et al. (2014) found evidence for a dually lateralized frontoparietal control network using the autonomy index. It was suggested that this control network acted as a coupler between the two hemispheres to increase efficiency while simultaneously supporting within-hemisphere processes. This was also evidenced by Spreng et al. (2013), which found that the frontoparietal control network exhibits distinct connectivity patterns with the default and attention networks in response to varying task requirements. Similarly, using a seed-based analysis, Braga et al. (2020) confirmed the presence of both left-lateralized and right-lateralized frontoparietal control networks. Collectively, these results point to control networks differentially executing cognitive processes within the left and right hemispheres.

Finally, the most right-lateralized network identified using NSAR was the Limbic-B network, a network that occupies cortical real estate associated with emotion (Olson et al., 2007; Pehrs et al., 2017; Sonkusare et al., 2020). Historically, emotion processing has been identified as being lateralized, perhaps beginning with lesion cases (Gainotti, 2019; Hughlings-Jackson, 1878; Luys, 1879). Later work suggested that specific aspects of emotion were lateralized, including the right-lateralization of emotion recognition, the right-lateralization of emotional control and expression, the right-lateralization of negative emotions, and the left-lateralization of positive emotions (Silberman & Weingartner, 1986). Contemporarily, it has been suggested that a hemispheric functional-equivalence hypothesis would better explain emotion neuroimaging results, such that emotion results from networks that are interrelated and may have different patterns of lateralization (for review, see Palomero-Gallagher & Amunts, 2022). This perspective emphasizes the intricate and interconnected nature of emotion-related neural processes, particularly those patterns of lateralization that emerge from inter-network relationships. Interestingly, the Limbic-B network appears to be at the center of our main results regarding lateralization relationships between networks.

### 4.3 Support for the Dependent Hypothesis of Network Lateralization

Beyond identifying networks with the greatest lateralization, we sought to understand how network lateralization was related between networks. Framing this investigation, a 2019 review described relationships between lateralized brain networks in terms of functional complementarity, or the degree of specialization minimizing functional overlap and redundancy for a pair of networks (Vingerhoets, 2019). Borrowing from the author’s ecological differentiation metaphor, just as species may specialize and fill different niches, facilitating coexistence among species, brain networks may also operate in a complementary fashion through the use of distinct computational processes and neural locations. Conversely, species (and brain networks) which do not specialize are functionally redundant and face increased competition. Crucially, this dynamic is such that high complementarity characterizes brain networks with low redundancy and competition while low complementarity describes brain networks with high redundancy and competition. To explore this further, we hypothesized that having one highly lateralized network corresponds with increased lateralization in other networks within an individual, and that this pattern of covariation would systematically occur across individuals (the dependent hypothesis). Interestingly, we identified lateralized network relationships exhibiting high and low complementarity occurring systematically across three different datasets.

#### 4.3.1 High Complementarity Network Relationships

Using correlation matrices, we found support for the dependent hypothesis in networks lateralized to contralateral hemispheres and exhibiting high complementarity. A negative relationship was found between the right-lateralized Limbic-B network and the left-lateralized Dorsal Attention-A network. Such a relationship is indicative of covariation, since negative NSAR values indicate left hemisphere lateralization, so greater lateralization of the left-lateralized Dorsal Attention-A network (negative NSAR values) were associated with greater lateralization of the right-lateralized Limbic-B network (positive NSAR values). Similarly, an additional negative relationship was identified between the right-lateralized Limbic-B network and the left-lateralized Default-C network. These relationships were systematic across individuals spanning three datasets, suggesting that there may be a population benefit to this configuration of lateralization. Interestingly, while others have suggested that the relationship between linguistic and spatial processing networks may be characterized by high complementary as well (Vingerhoets, 2019), we did not find evidence for this relationship in the present study.

#### 4.3.2 Low Complementarity Network Relationships

Beyond the high complementarity relationships, using correlation matrices we also identified a dependent relationship for networks lateralized to the same hemisphere exhibiting low complementarity. Since NSAR is derived based on surface area and the selected network parcellation method has a winner-takes-all approach, all cortical surface area for each individual is accounted for and networks lateralized to the same hemisphere are in competition with one another for cortical real estate. Thus, it is not surprising that two networks lateralized to the same hemisphere might have a negative relationship, such as that between the left-lateralized Dorsal Attention-A and Language networks identified in the present study. This relationship is such that as the lateralization for the Dorsal Attention-A network increases, the lateralization of the Language network decreases within individuals or vice versa, and this pattern was consistent across individuals from three datasets. Remarkably, two other examples of this low complementarity relationship have previously been identified and both involve language or linguistic processing. First, one group found evidence for the “co-lateralization” of language and praxis networks on both the individual and group levels (Vingerhoets et al., 2013). A similar “co-lateralization” relationship was identified for language and arithmetic regions (Pinel & Dehaene, 2010). In the latter study, it was suggested that “co-lateralization” might hint at the developmental effects of learning linguistic symbols on the organization of the arithmetic network. An additional hypothesis for this type of complementary relationship suggests that networks composed of overlapping nodes in the same hemisphere may be so similar that sharing proximate space is biologically less costly than the generation of a separate redundant network in the opposite hemisphere (Vingerhoets, 2019).

#### 4.3.3 Further Evidence of the Dependent Hypothesis

Additional support for the dependent hypothesis was found with the EFA and CFA structures across the three datasets. While we did not replicate the four-factor model from Liu et al. (2009), we did extract two factors for the HCP-Discovery dataset, which were then fitted in the HCP-Replication and HCPD datasets. Across these factor analyses, significant positive and negative loadings were found within each factor structure, suggesting that left-and right-lateralized networks work within a system level higher than the network.

#### 4.3.4 Characteristics of Lateralized Brain Network Organization

Together, the present evidence accumulated from the correlation matrices and EFA and CFA structures point to three overlapping features of organization for lateralized networks: complementarity, plasticity, and hierarchy. Beyond identifying which networks are lateralized, the present study evidences a configuration in which there are trade-offs in redundancy and competition. Rather than operating in isolation, lateralized networks appear to function in a larger system where their organization is interdependent. Given the zero-sum nature of a surface area-based approach, one might argue that this interconnectedness is an artifact. However, evidence from prior task fMRI lateralization research is in support of interconnectedness (Pinel & Dehaene, 2010; Vingerhoets et al., 2013). Similarly, the presented results demonstrating that network lateralization strength is related between networks suggests a degree of plasticity and adaptability in the brain’s functional organization. This potential developmental influence was hinted at with the “co-lateralization” of language and arithmetic; however, longitudinal models are needed to verify this speculation. Lastly, the present results describe a hierarchy of lateralized brain networks. This is most clearly demonstrated with the EFA and CFA structures, which were composed of both positive and negative factor loadings suggesting that these lateralized networks are not isolated but rather part of a larger system. This hierarchical organization of lateralized networks implies a mosaic of interaction and dependency within the broader brain architecture.

### 4.4 Limitations and Future Directions

One limitation to this work is that while functional connectivity may be constrained in part by anatomical connectivity, it is not necessarily dictated by anatomical connectivity. Several pieces of evidence point to this conclusion: functional connectivity is modulated by task (Shirer et al., 2012), recent experience (Lewis et al., 2009), caffeine (Laumann et al., 2015), and sleepiness (Tagliazucchi & Laufs, 2014); and is dynamic within a person over time (Hutchison et al., 2013). Furthermore, underlying brain geometry models of spontaneous neural activity appear to be more accurate and parsimonious than those derived from anatomical connectivity (Pang et al., 2023). Hence, NSAR as a connectivity and surface area-based measure is more reflective of functional rather than anatomical lateralization. As a result, future studies might benefit from exploring the Coutanche et al. (2023) method, which employs a surface-fingerprinting technique and multivariate laterality index for computing functional lateralization, offering a potentially complementary approach to NSAR in assessing functional lateralization.

In this study, individual parcellations were generated using the Kong et al. (2019) MS-HBM algorithm. However, improved versions of this algorithm have since been published (Kong et al., 2021; Yan et al., 2023), which account for parcel distributions, spatial contiguity, local gradients, and homotopy (or the lack thereof in Schaefer parcels). Thus, future investigations using NSAR might consider implementing an updated individual parcellation algorithm. Moreover, it would be valuable for future studies to explore lateralization in developmental and clinical populations to address questions regarding the developmental timeline of network lateralization and the potential disruptions in network lateralization observed in specific neurodevelopmental conditions such as autism or schizophrenia.

## 5 Conclusions

The present study investigated hemispheric asymmetries in the human brain, focusing on 17 functional networks. This was accomplished by implementing a surface area-based metric of lateralization, for which validity and reliability were examined. Following methodological development, we addressed two main questions: (1) which networks exhibit the greatest hemispheric asymmetries, and (2) how does lateralization in one network relate to the lateralization of other networks? We found that the Language, Dorsal Attention-A, and Default-C networks were significantly left-lateralized while the Visual-B, Salience/Ventral Attention-A, Control-B, Control-C, and Limbic-B networks were significantly right-lateralized. Additionally, using correlation matrices and EFA and CFA models to understand how lateralization is related between networks, we found general support for a dependent relationship between left-and right-lateralized networks. Within individuals, greater left-lateralization in a particular network (such as the Dorsal Attention-A or Default-C networks) was associated with greater right-lateralization in a particular network (such as the Limbic-B network). This pattern of lateralization appears to occur systematically across individuals, suggesting that lateralization follows a covariation paradigm. Further work is needed to understand how these findings may differ in developmental and clinical populations.

## 6 Acknowledgements

Data were provided in part by the Human Connectome Project, WU-Minn Consortium (principal Investigators: David Van Essen and Kamil Ugurbil; 1U54MH091657) funded by the 16 NIH Institutes and Centers that support the NIH Blueprint for Neuroscience Research; and by the McDonnell Center for Systems Neuroscience at Washington University. HCPD data reported in this publication was supported by the National Institute of Mental Health of the National Institutes of Health under Award Number U01MH109589 and by funds provided by the McDonnell Center for Systems Neuroscience at Washington University in St. Louis. The HCP-Development 2.0 Release data used in this report came from DOI: 10.15154/1520708. Collection of the NSD dataset was supported by NSF IIS-1822683 and NSF IIS-1822929. Furthermore, we acknowledge the support of the Office of Research Computing at Brigham Young University.

We are also grateful to Ru Kong for her assistance implementing the individual parcellation pipeline.

## 7 Data and Code Availability

With the exception of the HCPD dataset, the data reported on in the present study can be accessed publicly online (HCP: https://db.humanconnectome.org/; NSD: http://naturalscenesdataset.org/). The HCPD dataset is hosted through the NIMH Data Archive (NDA), through which access can be requested. Preprocessing and individual parcellation pipeline code are available through the CBIG repository on GitHub at https://github.com/ThomasYeoLab/CBIG. Scripts used to implement the processing pipelines and perform statistical analyses are available on GitHub at https://github.com/Nielsen-Brain-and-Behavior-Lab/NSAR2023.

## 9 Author Contributions/CRediT Statement

Madeline Peterson: Conceptualization, Methodology, Software, Validation, Formal analysis, Investigation, Data curation, Writing – original draft, Writing – review & editing, Visualization, and Project administration. Rodrigo M. Braga: Conceptualization, Writing – review & editing. Dorothea L. Floris: Writing – review & editing. Jared A. Nielsen: Conceptualization, Methodology, Writing — review and editing, Supervision, and Project administration.

