## Supplemental Figures, Tables, and Methods for "Evidence for a Compensatory Relationship between Left- and Right-Lateralized Brain Networks"

### Estimating Brain Network Asymmetries:

### The Development and Use of the Network Surface Area Ratio

Supplementary Materials

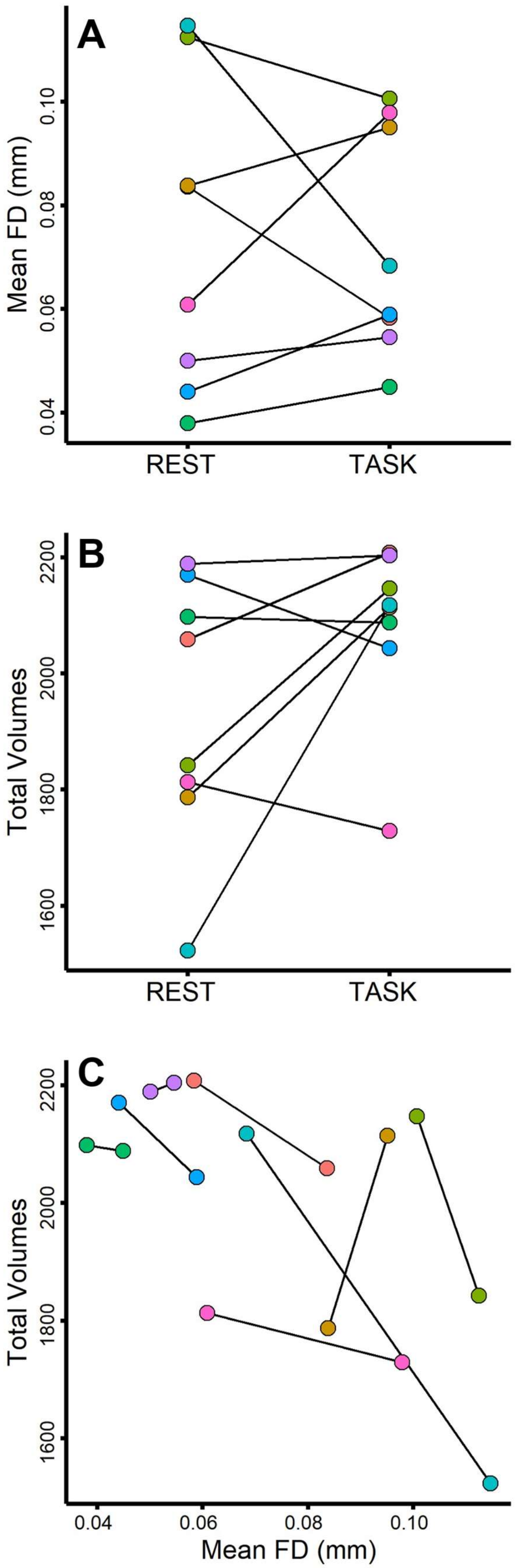
A
Mean FD (mm)
0.04
0.06
0.08
0.10
REST
TASK
B
Total Volumes
1600
1800
2000
2200
REST
TASK
C
Total Volumes
1600
1800
2000
2200
0.04
0.06
0.08
0.10
Mean FD (mm)

**Figure S1.** Data quality for the NSD dataset. Panel A depicts the mean framewise displacement across both resting-state and task fMRI data (12 runs each). Panel B depicts the total number of volumes post-motion correction available per participant for both resting-state and task fMRI data. Panel C depicts the total number of volumes available post-motion correction per participant by the mean framewise displacement. In each panel, a colored circle connected by a line to another colored circle represents the same individual.

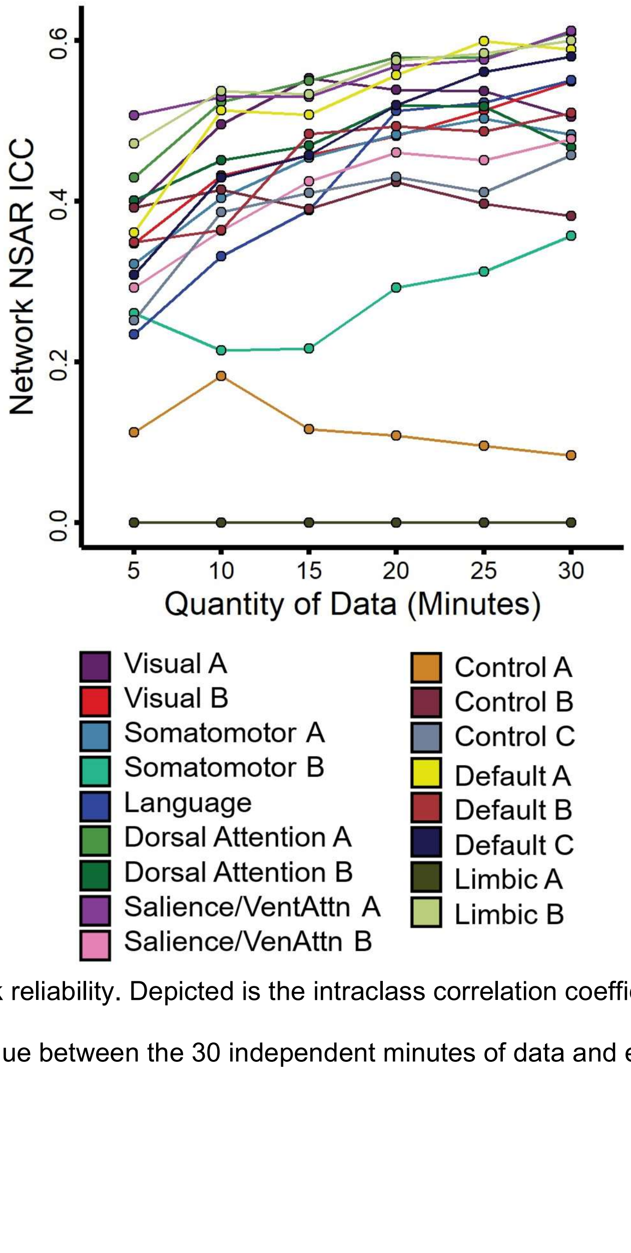


**Figure S2.** NSAR network reliability. Depicted is the intraclass correlation coefficient calculated for each network's mean NSAR value between the 30 independent minutes of data and each increment of data.

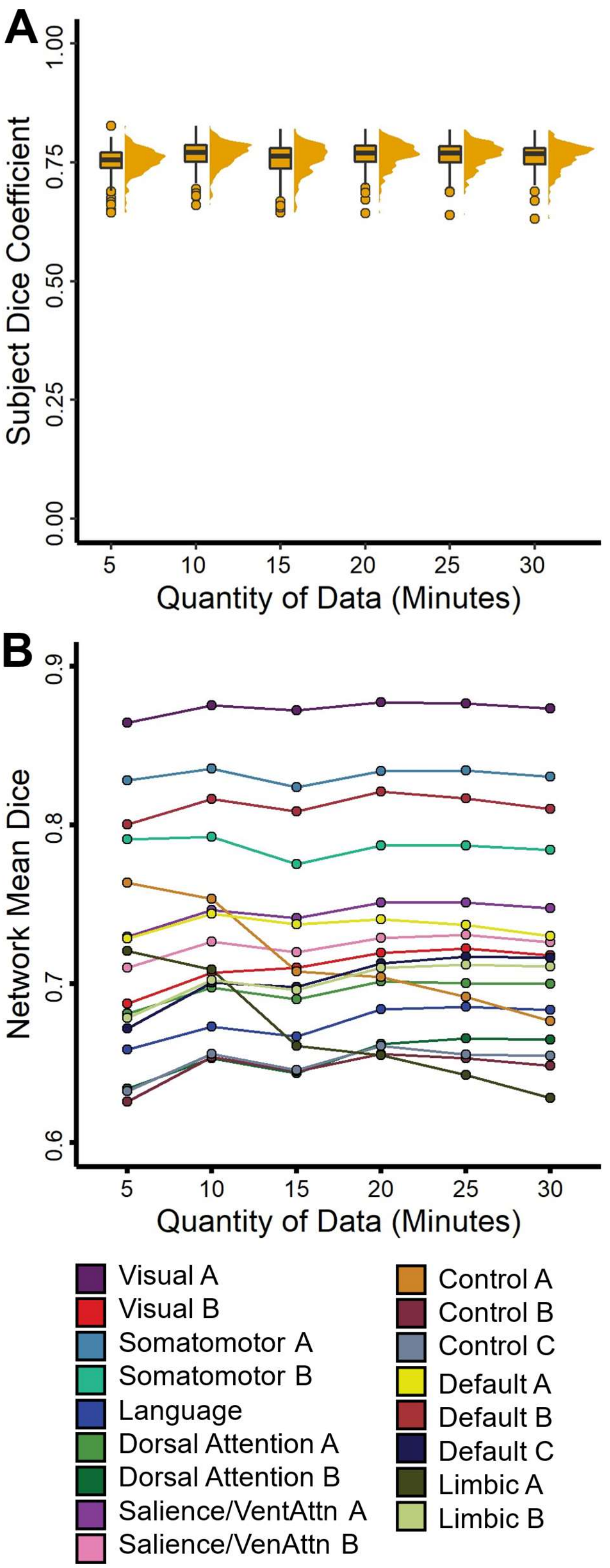


**Figure S3.** Parcellation overlap estimates in a subset of participants from the HCP dataset (*N* = 232).

Panel A depicts the subject dice coefficients calculated for the individual parcellations between each increment of data (5, 10, 15, … 30 minutes) and 30 independent minutes of data. Dice coefficient calculations are detailed in the Supplementary Methods. Comparable subject-level network assignment overlap analyses were conducted in the Midnight Scan Club dataset (see (Gordon et al., 2017) Figure 2 Panel B), finding that dice coefficients began at ~0.4 - 0.6 and plateaued at ~0.6 - 0.8. Panel B depicts the subject dice coefficients calculated for the individual parcellations between each increment of data (5, 10, 15, … 30 minutes) and 30 independent minutes of data averaged within each network. Comparable test-retest findings for parcellation overlap were described in Kong et al. (2019) Figures S10 Panel C and S11 Panel C, for which the somatomotor and visual networks exhibited the greatest intra-subject reproducibility.

**Supplementary Table 1**

*Identifying Specialized Networks Using Multiple Regressions in the HCP-Discovery (N = 276), HCP-Replication (N = 277), and HCPD (N = 343) Datasets*

| Network Intercept | Dataset | β | *SE* | *t* | *p* |
|---|---|---|---|---|---|
| **Visual-A** | | | | | |
| | HCP-DISC | 0.02 | 0.01 | 3.63 | < .001 |
| | HCP-REP | 0.01 | 0.01 | 2.79 | .006 |
| | HCPD | 0.01 | 0.00 | 1.54 | .13 |
| **Visual-B** | | | | | |
| | HCP-DISC | 0.07 | 0.02 | 3.95 | < .001 |
| | HCP-REP | 0.11 | 0.02 | 6.26 | < .001 |
| | HCPD | 0.11 | 0.01 | 8.18 | < .001 |
| Somatomotor-A | | | | | |
| | HCP-DISC | 0.02 | 0.01 | 2.92 | .004 |
| | HCP-REP | 0.01 | 0.01 | 1.41 | .16 |
| | HCPD | 0.01 | 0.01 | 1.22 | .22 |
| Somatomotor-B | | | | | |
| | HCP-DISC | -0.03 | 0.01 | -3.35 | < .001 |

| | | | | | |
|---|---|---|---|---|---|
| | HCP-REP | -0.03 | 0.01 | -2.85 | .005 |
| | HCPD | -0.01 | 0.01 | -3.42 | < .001 |
| **Language** | | | | | |
| | HCP-DISC | -0.15 | 0.03 | -4.71 | < .001 |
| | HCP-REP | -0.12 | 0.03 | -4.13 | < .001 |
| | HCPD | -0.19 | 0.02 | -8.11 | < .001 |
| **Dorsal Attention-A** | | | | | |
| | HCP-DISC | -0.25 | 0.02 | -12.05 | < .001 |
| | HCP-REP | -0.31 | 0.02 | -15.37 | < .001 |
| | HCPD | -0.22 | 0.02 | -13.74 | < .001 |
| Dorsal Attention-B | | | | | |
| | HCP-DISC | -0.02 | 0.02 | -1.24 | .21 |
| | HCP-REP | -0.03 | 0.02 | -1.47 | .14 |
| | HCPD | 0.01 | 0.01 | 0.87 | .38 |
| **Salience/VenAttn-A** | | | | | |
| | HCP-DISC | 0.05 | 0.01 | 4.48 | < .001 |
| | HCP-REP | 0.07 | 0.01 | 6.05 | < .001 |
| | HCPD | 0.05 | 0.01 | 6.35 | < .001 |
| Salience/VenAttn-B | | | | | |
| | HCP-DISC | -0.02 | 0.01 | -1.59 | .11 |
| | HCP-REP | -0.03 | 0.01 | -2.56 | .01 |
| | HCPD | -0.03 | 0.01 | -3.52 | < .001 |
| **Control-A** | | | | | |
| | HCP-DISC | -0.07 | 0.02 | -3.7 | < .001 |
| | HCP-REP | -0.11 | 0.02 | -6.51 | < .001 |
| | HCPD | -0.05 | 0.01 | -6.52 | < .001 |
| **Control-B** | | | | | |
| | HCP-DISC | 0.19 | 0.02 | 7.95 | < .001 |

| | | | | | |
|---|---|---|---|---|---|
| | HCP-REP | 0.18 | 0.02 | 8.09 | < .001 |
| | HCPD | 0.14 | 0.01 | 9.93 | < .001 |
| **Control-C** | | | | | |
| | HCP-DISC | 0.09 | 0.02 | 6.11 | < .001 |
| | HCP-REP | 0.09 | 0.01 | 7.16 | < .001 |
| | HCPD | 0.09 | 0.01 | 9.53 | < .001 |
| Default-A | | | | | |
| | HCP-DISC | -0.02 | 0.02 | -1.01 | .31 |
| | HCP-REP | -0.02 | 0.02 | -1.52 | .13 |
| | HCPD | -0.01 | 0.01 | -1.04 | .3 |
| Default-B | | | | | |
| | HCP-DISC | 0.01 | 0.01 | 1.04 | .29 |
| | HCP-REP | 0.02 | 0.01 | 2.13 | .03 |
| | HCPD | 0.01 | 0.01 | 0.87 | .39 |
| **Default-C** | | | | | |
| | HCP-DISC | -0.21 | 0.02 | -10.56 | < .001 |
| | HCP-REP | -0.17 | 0.02 | -9.37 | < .001 |
| | HCPD | -0.15 | 0.02 | -8.83 | < .001 |
| Limbic-A | | | | | |
| | HCP-DISC | 0.06 | 0.01 | 4.4 | < .001 |
| | HCP-REP | 0.09 | 0.01 | 6.3 | < .001 |
| | HCPD | 0.01 | 0.01 | 0.91 | .37 |
| **Limbic-B** | | | | | |
| | HCP-DISC | 0.25 | 0.03 | 8.93 | < .001 |
| | HCP-REP | 0.32 | 0.03 | 11.89 | < .001 |
| | HCPD | 0.28 | 0.02 | 12.21 | < .001 |

*Note:* Coefficients and *p*-values for the intercept are shown. None of the covariates (mean-centered age, mean-centered framewise displacement, handedness, and sex) were consistently significant across the three datasets for any of the networks. Networks with reliably significant (Bonferroni-corrected alpha level of .003) intercepts are bolded.

**Supplementary Table 2**

*Left-lateralized Network Comparisons*

| Network Comparison | Dataset | β | *SE* | *t* | *p* |
|---|---|---|---|---|---|
| Language | | | | | |
| Dorsal Attention-A | | | | | |
| | HCP-DISC | -0.11 | 0.02 | -6.98 | < .001 |
| | HCP-REP | -0.11 | 0.02 | -6.73 | < .001 |
| | HCPD | -0.12 | 0.01 | -9.69 | < .001 |
| Default-C | | | | | |
| | HCP-DISC | 0.09 | 0.01 | 7.94 | < .001 |
| | HCP-REP | 0.1 | 0.01 | 8.63 | < .001 |
| | HCPD | 0.11 | 0.01 | 9.09 | < .001 |
| Dorsal Attention-A | | | | | |
| Default-C | | | | | |
| | HCP-DISC | -0.01 | 0.02 | -0.93 | .36 |
| | HCP-REP | -0.00 | 0.02 | -0.31 | .76 |
| | HCPD | -.02 | 0.01 | -1.29 | .19 |

*Note:* Comparisons consist of multiple regressions, which included a network variable with two levels (the two networks under comparison), mean-centered age, sex, handedness, and mean-centered mean framewise displacement.

**Supplementary Table 3**

*Right-lateralized Network Comparisons*

| Network Comparison | Dataset | *β* | *SE* | *t* | *p* |
|---|---|---|---|---|---|
| Visual-B | | | | | |
| Salience/VenAttn-A | | | | | |
| | HCP-DISC | -0.04 | 0.01 | -4.56 | < .001 |
| | HCP-REP | -0.05 | 0.01 | -5.04 | < .001 |
| | HCPD | -0.05 | 0.01 | -6.69 | < .001 |
| Control-B | | | | | |
| | HCP-DISC | 0.09 | 0.02 | 5.69 | < .001 |
| | HCP-REP | 0.08 | 0.01 | 6.19 | < .001 |
| | HCPD | 0.05 | 0.01 | 5.47 | < .001 |
| Control-C | | | | | |
| | HCP-DISC | -0.01 | 0.01 | -1.03 | .3 |
| | HCP-REP | -0.01 | 0.01 | -1.27 | .2 |

| | | | | | |
|---|---|---|---|---|---|
| | HCPD | -0.01 | 0.01 | -1.56 | .12 |
| Limbic-B | | | | | |
| | HCP-DISC | 0.08 | 0.02 | 4.21 | < .001 |
| | HCP-REP | 0.18 | 0.01 | 12.89 | < .001 |
| | HCPD | 0.18 | 0.01 | 13.46 | < .001 |
| Salience/VenAttn-A | | | | | |
| Control-B | | | | | |
| | HCP-DISC | 0.06 | 0.01 | 4.32 | < .001 |
| | HCP-REP | 0.12 | 0.01 | 11.18 | < .001 |
| | HCPD | 0.11 | 0.01 | 13.1 | < .001 |
| Control-C | | | | | |
| | HCP-DISC | 0.03 | 0.01 | 3.71 | < .001 |
| | HCP-REP | 0.03 | 0.01 | 4.29 | < .001 |
| | HCPD | 0.04 | 0.07 | 6.04 | < .001 |
| Limbic-B | | | | | |
| | HCP-DISC | 0.21 | 0.01 | 16.23 | < .001 |
| | HCP-REP | 0.22 | 0.01 | 17.81 | < .001 |
| | HCPD | 0.23 | 0.01 | 19.23 | < .001 |
| Control-B | | | | | |
| Control-C | | | | | |
| | HCP-DISC | -0.09 | 0.01 | -6.97 | < .001 |
| | HCP-REP | -0.09 | 0.01 | -7.78 | < .001 |
| | HCPD | -0.07 | 0.01 | -7.59 | < .001 |
| Limbic-B | | | | | |
| | HCP-DISC | 0.09 | 0.02 | 5.89 | < .001 |
| | HCP-REP | 0.1 | 0.02 | 6.74 | < .001 |
| | HCPD | 0.12 | 0.01 | 9.32 | < .001 |
| Control-C | | | | | |
| Limbic-B | | | | | |
| | HCP-DISC | 0.18 | 0.01 | 13.04 | < .001 |
| | HCP-REP | 0.19 | 0.01 | 14.69 | < .001 |
| | HCPD | 0.19 | 0.01 | 15.35 | < .001 |

*Note:* Comparisons consist of multiple regressions, which included a network variable with two levels (the two networks under comparison), mean-centered age, sex, handedness, and mean-centered mean framewise displacement.

#### Supplementary Methods

This section provides additional information on estimating the quantity of data needed to generate stable individual parcellation labels, similar to the in-text analysis performed with NSAR values.

##### Stable Estimate Analysis for Individual Parcellation Labels

A dice coefficient (Dice, 1945; Sorenson, 1948) was calculated in order to identify parcellation label overlap between the parcellations resulting from iteration (5, 10, 15, etc. minutes) and the parcellation resulting from the independent 30 minutes of data. The dice coefficient is calculated as follows:

$$\text{Dice} = \frac{2|X \cap Y|}{|X|+|Y|}$$

where $X \cap Y$ represents the number of vertices with the same network labels in the same positions across the iteration parcellation and the independent 30 minutes parcellation. The denominator represents the total number of vertices with a given network label across both the iteration parcellation and the independent 30 minutes parcellation. Dice coefficients were also estimated for each network within each participant between each increment of data and the 30 independent minutes of data, and then an average dice coefficient for each network was computed for each increment of data.

When a single dice coefficient was calculated within individuals, only five minutes of data were needed obtain a mean dice coefficient of 0.75 (*SD* = 0.03), which was highly similar to the 0.76 mean (*SD* = 0.03) dice coefficient obtained with 30 minutes of data (see Supplementary Figure S3 Panel A). Networks with the highest mean dice coefficients include those related to sensory and somatomotor functions such as the Visual-A, Visual-B, Somatomotor-A, and Somatomotor-B networks (see Supplementary Figure S3 Panel B).
